# Asymmetry-induced distinct mechanisms and the transporting role of sodium in bacterial fluoride channel Fluc

**DOI:** 10.64898/2026.02.18.706662

**Authors:** Fernando Montalvillo Ortega, Kira Mills, Hedieh Torabifard

## Abstract

Bacterial fluoride channels (Fluc) are exceptional in their ability to conduct fluoride ions at rapid rates while maintaining extraordinary selectivity, an unusual property among anion transporters. Fluc achieves this through an antiparallel homodimeric assembly that forms two asymmetric ion-conduction pores and hosts a centrally bound sodium ion of previously uncertain function. Despite extensive structural characterization, the molecular basis for Fluc’s dual-pore asymmetry, transport efficiency, and sodium involvement has remained unresolved. Here, we performed long-timescale molecular dynamics simulations under electrophysiological conditions to elucidate the mechanisms of fluoride translocation through each pore. Our results reveal two distinct conduction modes: Pore I operates via the Channsporter mechanism, characterized by single-ion transport; while Pore II follows the Multi-ion mechanism in which paired fluoride ions exploit electrostatic repulsion to achieve faster sequential translocations. These findings demonstrate that structural asymmetry gives rise to mechanistic specialization within the same homodimeric protein. We further propose that the central sodium ion serves as a dynamic cofactor in fluoride transport, coupling its vertical oscillation to fluoride movement in a pore-dependent manner. Together, our results establish a unified model of asymmetric dual-pore conduction in Fluc and introduce a broader paradigm in membrane transport biology, one where inherent structural asymmetry enables divergent yet coexisting mechanisms of ion conduction.

**Significance:** Fluoride is a naturally occurring environmental ion that is toxic to bacteria at elevated concentrations, necessitating specialized export systems for survival. Bacteria encode two distinct fluoride exporters: the CLC^*F*^ antiporter and the Fluc channel. The latter of which remains mechanistically unresolved due to its unusual dual-pore architecture and lack of close homologs. Using extensive molecular dynamics simulations, we dissect fluoride permeation through Fluc and show that its two antiparallel pores operate through distinct transport mechanisms. Besides, we uncover a previously unrecognized role for the central sodium ion as an essential cofactor that enables fluoride conduction. Together, these findings provide new mechanistic insight into fluoride export and clarify the functional asymmetry underlying Fluc-mediated transport.

---

Fluorine is among the most abundant elements in the Earth’s crust, yet, because of its great reactivity, it exists as inorganic fluorides rather than in its elemental form (1, 2). Environmental fluoride (F−) is mobilized into soils and waters through weathering and erosion of fluoride-bearing minerals, volcanic eruptions, and human intervention (2, 3). These processes have led to an increase in the bioavailability of this anion, which has toxic effects at high concentrations. F^−^ acts as an antimicrobial agent by inhibiting essential metabolic enzymes, displacing native electronegative substrate groups (4–6). Despite this latter fact, for many years, the biological relevance of fluoride transport across membranes was largely overlooked (7). Apart from passive intracellular accumulation via ion-trapping of its conjugate acid HF (8, 9), which readily permeates membranes at rates comparable to water, fluoride membrane biology was considered negligible (10).

This paradigm shifted with the discovery of fluoride-responsive riboswitches in bacteria by Baker *et al*. (11). The crcB RNA motif was identified as a fluoride-sensing riboswitch that regulates the expression of two distinct classes of bacterial fluoride ion exporters: CLC^*F*^ F^−^/H^+^ antiporters (12) and fluoride channels Fluc (13). Concurrently, it was recognized that eukaryotic organisms also rely on dedicated fluoride export as mediated by the Fluc homolog FEX (fluoride exporter) (14). While CLC^*F*^ proteins, despite being phylogenetically distant, bear mechanistic similarities to canonical CLC transporters (12, 15), Fluc and FEX represent a unique and enigmatic system with no structural homologs (13, 16).

To date, Fluc remains the most selective ion transporter reported (10, 13, 17). Remarkably, its rapid conductivity with extraordinarily high selectivity arises from an equally unusual architecture (10, 13, 17, 18). As illustrated in Fig. 1A–B, Fluc forms an antiparallel homodimer, a rare arrangement with only a handful of known examples (19, 20). Furthermore, unlike typical channel assemblies that generate a single central pore, Fluc creates two independent lateral pores (Fig. 1B), a feature confirmed through structural and functional studies (7, 17, 18). Owing to this antiparallel configuration, each pore is intrinsically asymmetric. Additionally, as displayed in Fig. 1C-D, Fluc proteins present a central sodium ion (Na^+^) bound within the protein structure whose structural or mechanistic role remains unresolved (21, 22). Although some transporters that cotransport sodium ions have a similar binding nature to the sodium ion in Fluc, Fluc is the exceptional case of a sodium ion utilized for structural or mechanistic purposes (21, 23).

**Fig. 1.**
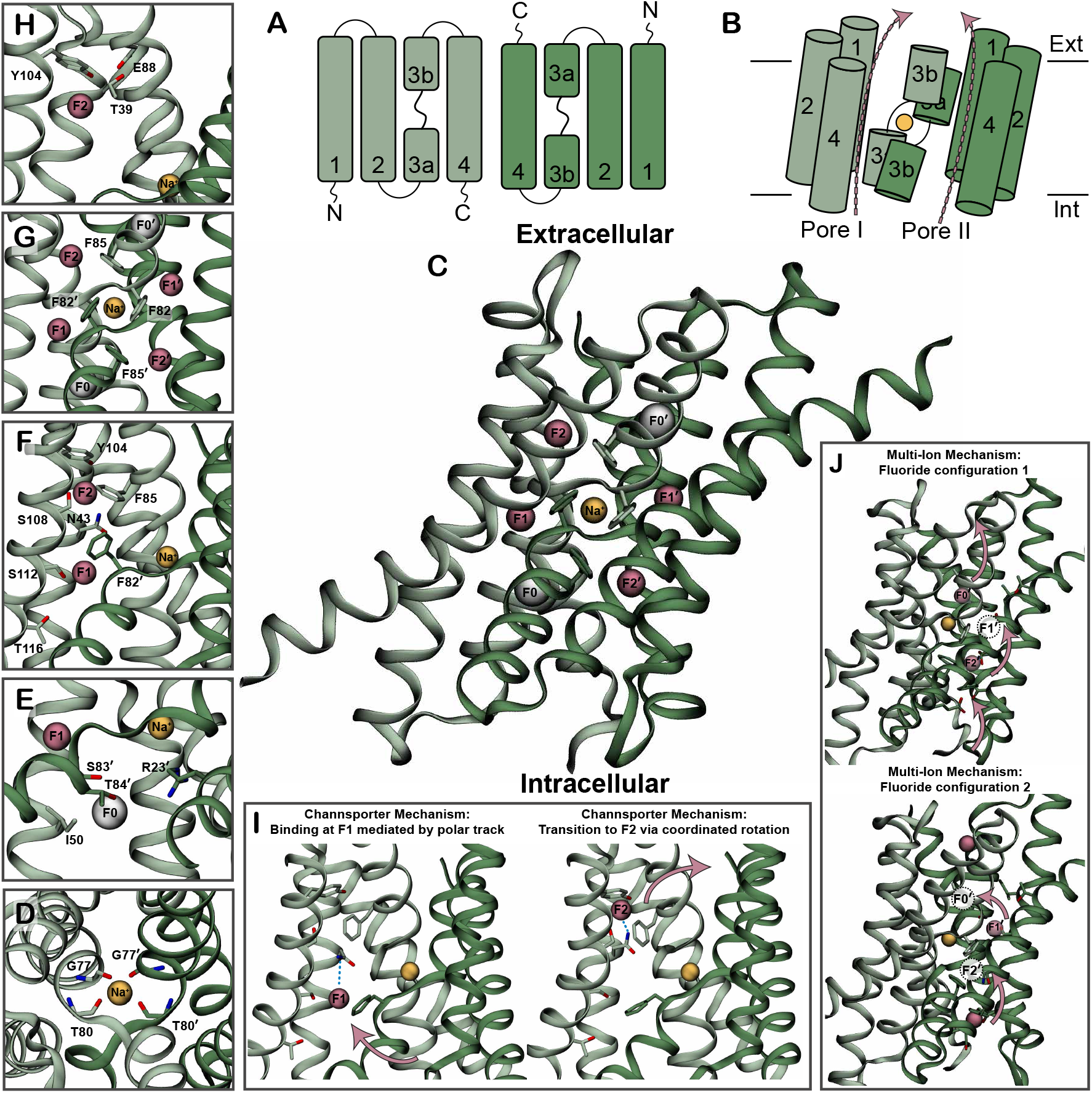
Summary of Fluc’s critical structural and mechanistic features. (*A*) Topology of Fluc highlighting its antiparallel homodimeric assembly with labeled helices. (*B*) Schematic representation of the Fluc homodimer assembly, with pink arrows indicating the two pore pathways and the central sodium shown as a yellow sphere. (*C*) Crystal structure of Fluc-*Bpe* with fluoride, bromide, and sodium crystallized positions highlighted as pink, white, and yellow spheres, respectively. (*D*) Top-view zoom-in of the central sodium binding site. (*E*) Close-up view of the F0 binding site. (*F*) Close-up view of the polar track containing F1 and F2 binding sites. (*G*) Close-up view of the Phe-box. (*H*) Close-up view of the anion-recognition TEY triad. (*I*) Schematic of the proposed Channsporter mechanism. (*J*) Schematic of the proposed Multi-ion mechanism.

Substantial efforts have been devoted to elucidating the mechanism of fluoride permeation in Fluc (7, 17, 21, 22, 24– 27). Crystallographic studies with fluoride and bromide (Br^−^) identified three discrete anion binding sites (17, 24), designated F0, F1, and F2 (Fig. 1C). A closer view of F0 bromide binding site provided in Fig. 1E, reveals a binding mode supported by hydrogen-bonding residues. Mutagenesis of residues surrounding F0 severely impaired transport but had no effect on selectivity (24). Simulation studies further demonstrated that replacing Br^−^ with F^−^ allows the ion to migrate closer to the central sodium, approaching within ∼4.5 Å (25).

From F0, the ion progresses into the pore through binding sites F1 and F2, zoomed in Fig. 1F. These sites are stabilized by the polar track, formed by conserved hydrogen-bond donors along transmembrane helix 4, the highly conserved N43 of helix 2, and the aromatic side chains of the Phe-box, shown in Fig. 1G. Collectively, these motifs establish a coordinated pathway that guides fluoride through the channel interior (7, 17, 26). The functional importance of the polar track residues, however, appears to vary among homologs. For instance, F82I in Fluc-*Bpe* is strongly detrimental, whereas F85I is tolerated (17); in Fluc-*Ec2*, F80I and F83I render the protein inactive,(7) while in the eukaryotic FEX1 from *Saccharomyces cerevisiae*, F96A and F330A only reduce transport modestly (28). Strikingly, simultaneous mutation of the two conserved phenylalanines consistently abolishes activity across homologs (7, 28), suggesting that fluoride coordination along the pore depends on a complex and interdependent network of interactions.

Finally, to complete translocation, fluoride must traverse the TEY triad, depicted in Fig. 1H. Mutations at this location usually result in anemic transports of fluoride, highlighting their importance (24). Notably, in the opposing pore of the antiparallel dimer, ions encounter these features in reverse order: entering first through the TEY triad, which has been proposed by McIlwain *et al*. to serve as an anion recognition motif and selectivity gate (24), then passing the polar track and F1–F2 sites, and ultimately exiting via F0 into the electropositive vestibule.

Despite these structural insights, the precise transport mechanism and basis of selectivity remain unresolved. Two hypotheses dominate: (i) the Channsporter mechanism, in which a single ion translocates via coordinated rotations of hydrogen-bonding residues (17), depicted in Fig. 1I, and (ii) the Multi-ion mechanism, in which ions move cooperatively, propelled by electrostatic repulsion (24, 25), showcased in Fig. 1J. In both models, selectivity is thought to arise from complete dehydration of fluoride, as observed in other ion channels (17, 24, 27). Yet, mutagenesis studies have failed to conclusively validate either mechanism or alter selectivity. The central sodium ion further complicates the picture. While McIlwain *et al*. demonstrated that replacing sodium with lithium (Li^+^) abolishes transport (21), despite lithium’s seemingly more favorable coordination, its precise mechanistic role remains obscure. In support of this, Kang *et al*. confirmed sodium-dependent transport and its abolition upon lithium substitution in the eukaryotic homolog FEX (23).

To address these outstanding questions, we performed long-timescale molecular dynamics (MD) simulations under electrophysiological conditions. Our double-bilayer system recreated electrochemical gradients experienced *in vivo*, allowing unbiased sampling of conduction events. Strikingly, we observed distinct behaviors in the two pores: one consistent with the Channsporter mechanism, the other with the Multi-ion mechanism. Besides, we identified sodium ion displacements along the bilayer normal that underlie a mechanistic role in fluoride transport. Collectively, our results provide new perspectives on the architecture, mechanism, and selectivity of Fluc channels, offering hypotheses for future experimental validation. More broadly, our findings highlight a novel paradigm in membrane transport biology: channels capable of sustaining dual, asymmetric pores within a single homodimeric framework, aided by a central sodium ion functioning as a cofactor in transport.

## Results

### CompEL Simulations to Study Rapid Fluoride Transport in Fluc

Computational electrophysiology (CompEL) simulation protocols have become a valuable tool for studying ion channel transport due to their ability to model translocation events under applied electrochemical gradients without introducing artificial forces or external potentials (29, 30). Fluc exhibits exceptionally fast fluoride transport, with approximately 10^6^ ion translocations per second (10, 26, 31), corresponding to roughly one translocation per microsecond. These timescales are accessible with current computational resources, making Fluc an ideal system for long-timescale, unbiased MD simulations.

To investigate fluoride permeation in fluoride channels, we employed a dual-membrane electrophysiology setup capable of supporting electrochemical gradients across the protein-containing membrane. This configuration is essential for overcoming the constraints imposed by periodic boundary conditions (PBCs), which otherwise prevent the establishment of net transmembrane gradients. To enhance the likelihood of observing spontaneous ion translocation events during the simulations, we systematically varied the electrochemical driving forces by combining multiple intracellular KF concentrations (0.15 M, 0.25 M, and 0.5 M) with charge imbalances of 0e, 4e, and 8e. A detailed overview of these conditions is provided in the Methods section and in Fig. S1A and Table S1.

Additionally, given the ongoing debate regarding Fluc’s fluoride crystallized locations (22, 25), we explored how different initial fluoride-binding configurations might be favored. Six initial occupation states were tested, including fully occupied, fully empty, one pore occupied, intracellularly occupied, extracellularly occupied, and diagonally occupied, summarized in Fig. S1B. These configurations were to assess whether certain ion distributions are energetically favored or promote translocation.

### Initial Observations Reveal Pores with Distinct Natures

The total simulation time amounted to 108 *µ*s, comprising 54 independent 2 *µ*s trajectories with distinct binding site configurations, intracellular KF concentrations, and charge imbalances. To ensure proper equilibration, the first 100 ns of each production were excluded from analysis based on the stabilization of the backbone RMSD shown in Fig. S2A. The accuracy of our simulations is supported by the RMSF results presented in Fig. S2B-C that align closely with the solid-state NMR data reported by Zhang *et al*., identifying three highly flexible regions spanning residues 25–35, 50–70, and 88–103 (25).

During equilibration, fluoride ions were unrestrained, allowing spontaneous redistribution from their initially assigned binding site, and when energetically favorable, escape from the protein entirely to reach more stable configurations. To monitor fluoride dynamics and translocation events, both trajectory visualization and in-house Tcl and Python scripts were employed to quantify the number of fluoride ions residing within the Fluc homodimer and in the extracellular and intracellular compartments, with results per trial displayed in Fig. S3. As presented in Fig. 2A-B, the script used a set of orthogonal planes to accurately track fluoride positions along the bilayer normal, dividing the system into six compartments: intracellular, Pore I lower region (F0-F1), Pore II lower region (F2^*′*^), Pore I upper region (F2), Pore II upper region (F0^*′*^-F1^*′*^), and extracellular. Additional compartments that would distinguish between the crystallographic F0 and F1 sites were not utilized due to inability to reliably differentiate between these positions in our simulations. Consistent with the observations of Zhang *et al*. (25), fluoride ions rarely occupied the crystallographically assigned F0 and F1 sites as reported in Fig. 2C. Instead, F^−^ remained within approximately 5 Å of the central sodium ion, preferentially sampling a position best described as a *quasi*-F0 site. This observation suggests that the F0 and F1 sites may represent nonphysiological crystallographic artifacts rather than functionally relevant binding positions. Accordingly, the six-region division defined here offers a more accurate and easily interpretable framework for describing fluoride occupancy within the Fluc pores.

**Fig. 2.**
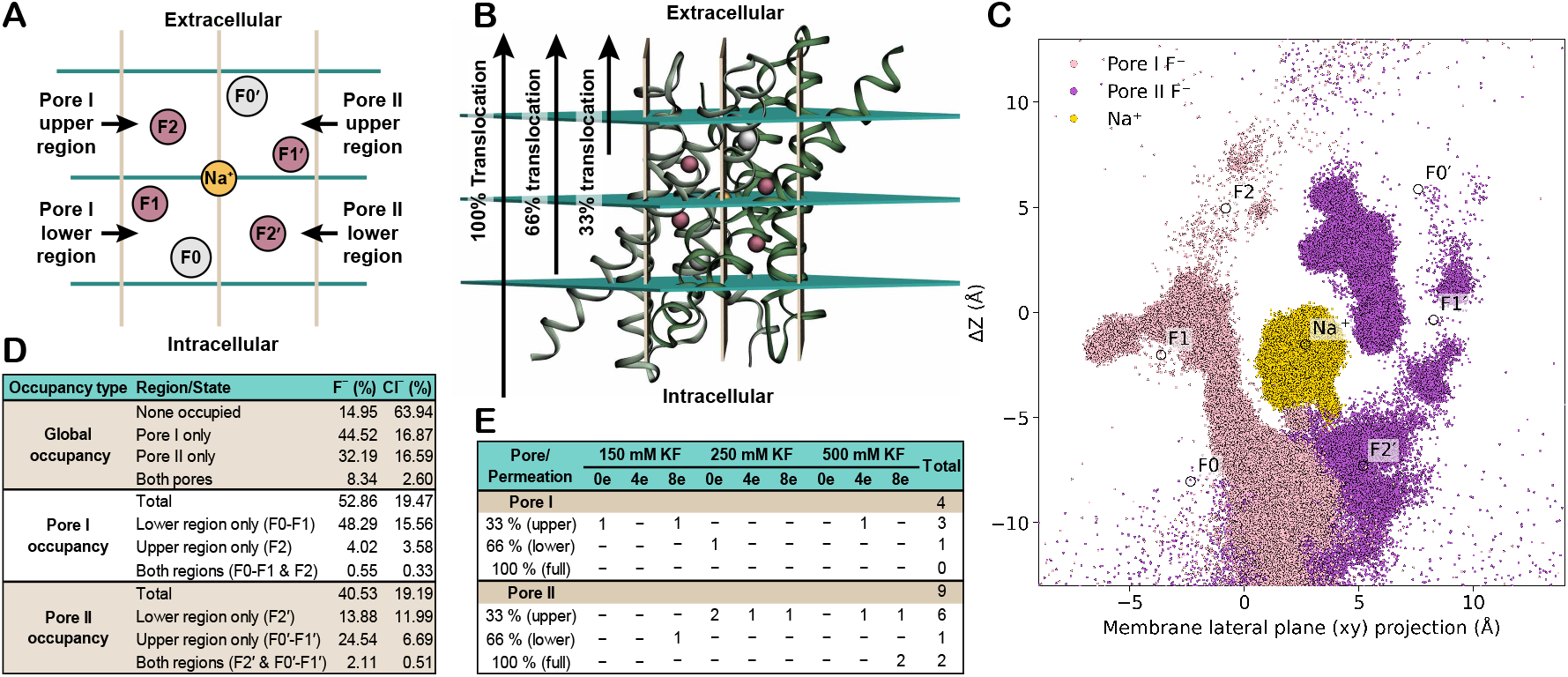
Quantification of pore occupancy and fluoride translocation in Fluc. (*A*) 2D Schematic representation of the six compartments delineated thanks to the orthogonal planes strategy: the intracellular, Pore I lower region (F0 and F1), Pore II lower region (F2^*′*^), Pore I upper region (F2), Pore II upper region (F1^*′*^ and F0^*′*^), and extracellular compartments. 3D Schematic representation of the orthogonal planes described in panel *A* used to determine pore occupancy and translocation events. Three translocation types were defined according to the initial fluoride position after a 100 ns equilibration period: 100 % translocation if the fluoride originated in the intracellular region, 66 % if it started in a lower-pore region, and 33 % if it began in an upper-pore region. (*C*) Scatter plot illustrating the motion of the central sodium ion (yellow dots) alongside with Pore I (pink) and Pore II (purple) assigned fluoride ions. The plot highlights the high mobility of sodium and delineates the conductive pathways of fluoride through each pore. (*D*) Global, pore-specific, and region-partitioned occupancy for anions across all simulations. (*E*) Fluoride permeation events categorized by extent of pore traversal across all simulations.

Analysis of fluoride occupancy after equilibration, presented in Fig. 2D, revealed that the regions containing the crystallographic F0 and F1 sites, namely the Pore I lower region and Pore II upper region, were the most frequently populated, accounting for 48.84% of the simulation time in Pore I and 26.65% in Pore II. In contrast, the polar track regions, predominantly represented by the Pore I upper region and Pore II lower region, were rapidly vacated, exhibiting lower occupancies of 4.57% and 15.99%, respectively (Fig. 2D). To probe the mechanistic question of whether transport occurs via a single-ion (“Channsporter”) or Multi-ion mechanism, we further quantified double-occupancy events, defined as simultaneous fluoride presence in both the upper and lower regions of a given pore. Strikingly, double occupancy was rare in Pore I, occurring in only 0.55% of all simulated time, whereas Pore II exhibited a considerably higher rate of 2.11% (Fig. 2D). This is approximately a four-fold increase that suggests distinct ion-handling behaviors between the two pores.

This divergence became even more pronounced when examining fluoride translocation events as shown in Fig. 2E. Full translocations were exceedingly rare, underscoring the inherent difficulty of capturing complete permeation events in narrow channels such as Fluc, even under long-timescale simulations and varying electrochemical conditions. Nevertheless, one trajectory displayed two complete translocations through Pore II, whereas no full translocation events were observed in Pore I. Partial translocations were also uncommon: two 66% events (one per pore) and nine 33% events (three in Pore I and six in Pore II) were detected (Fig. 2E). In summary, a total of thirteen distinct permeation events occurred, four through Pore I and nine through Pore II (Fig. 2E). These results indicate that under the simulated conditions, Pore II exhibits a higher fluoride-transport efficiency than Pore I.

To probe the basis of Fluc selectivity, the same approach was also applied to track chloride counter-ion dynamics. As reported in Fig. 2D, and in contrast to fluoride, chloride ions predominantly occupied the lower pore regions near the intracellular side and failed to progress further into the channels, resulting in no translocation events. This behavior is noteworthy, as it suggests that the electrochemical gradient applied during the simulations may enhance anion sampling at the pores entry, as expected. Nevertheless, most simulations contained higher fluoride than chloride concentrations, potentially introducing bias when directly comparing their behaviors. To minimize this, we examined anion occupancies only under conditions in which both ions were present at equal concentrations (150 mM KF), and under equivalent conditions with initially unoccupied binding sites (150 mM KF and Empty), as reported in Table S2. Across all scenarios, the outcome remained consistent: fluoride ions sampled intracellular binding sites significantly more frequently than chloride, indicating that the electropositive vestibule contributes to the early stages of anion selectivity.

Within this vestibule, we observed a particularly striking behavior of Arg23^*′*^ as displayed in Fig. S4 and Movie S1. The residue appears to actively recruit fluoride from the intracellular medium (Fig. S4A) and distribute it into both pores (Fig. S4B), effectively committing each fluoride to a specific conduction pathway. This interpretation is supported by the z-coordinate distributions and frequency of salt-bridge interactions between fluoride and Arg23^*′*^ (Fig. S4C–F).

### Pore I Uses the Channsporter Mechanism for Fluoride Conduction

Given the distinct behaviors exhibited by the two pores in double-region occupancy, translocation efficiency, and arginine-guided fluoride distribution, all subsequent analyses were conducted separately for each pore. Using the results from the orthogonal-plane classification (Fig. 2), fluoride ions interacting with Fluc were assigned to either Pore I or Pore II based on where they spent more than 50% of their interaction time with the protein (Fig. 2C). Once classified, the fluoride trajectories were analyzed to identify key physicochemical factors underlying their conduction mechanism and contribution to Fluc’s selectivity in each pore.

Analysis of Pore I began by examining the hydration state of fluoride ions along the bilayer normal as displayed in Fig. 3A. Consistent with the findings of Yue *et al*. (27), fluoride underwent progressive dehydration as it penetrated deeper into the polar track, reaching a minimum hydration level of one to two water molecules near the center of the Pore I lower region (F0 and F1 containing compartment). The substantial energy penalty associated with fluoride dehydration (32) was counterbalanced by strong protein–ion interactions, as reflected in the fluoride-protein Linear Interaction Energy (lie) profile in Fig. 3A. The correlation between lie and the number of hydrogen bonds, shown in Fig. 3A, suggests that much of this energetic compensation arises from hydrogen bonding with the protein. The maximum hydrogen-bond count of approximately four to five occurred near the center of the Pore I lower region, coinciding with the zone of minimal hydration. As fluoride progressed toward F2, some hydrogen bonds were replaced by additional water molecules or by edge-on anion–*π* interactions with phenylalanine residues of the Phe-box, as showcased in Fig. 3B. These edge-on anion–*π* interaction profiles are consistent with the original work on Fluc-*Bpe* by Stockbridge *et al*., in which the F82I mutation was found to be strongly detrimental to transport, whereas F85I had a lesser effect (17). This is in agreement with our simulations, which reveal a persistent edge-on anion–*π* interaction involving F82^*′*^. Inspection of the trajectories confirmed that all conduction events through Pore I involved single fluoride ions, supporting a Channsporter-type mechanism, see Movie S2.

**Fig. 3.**
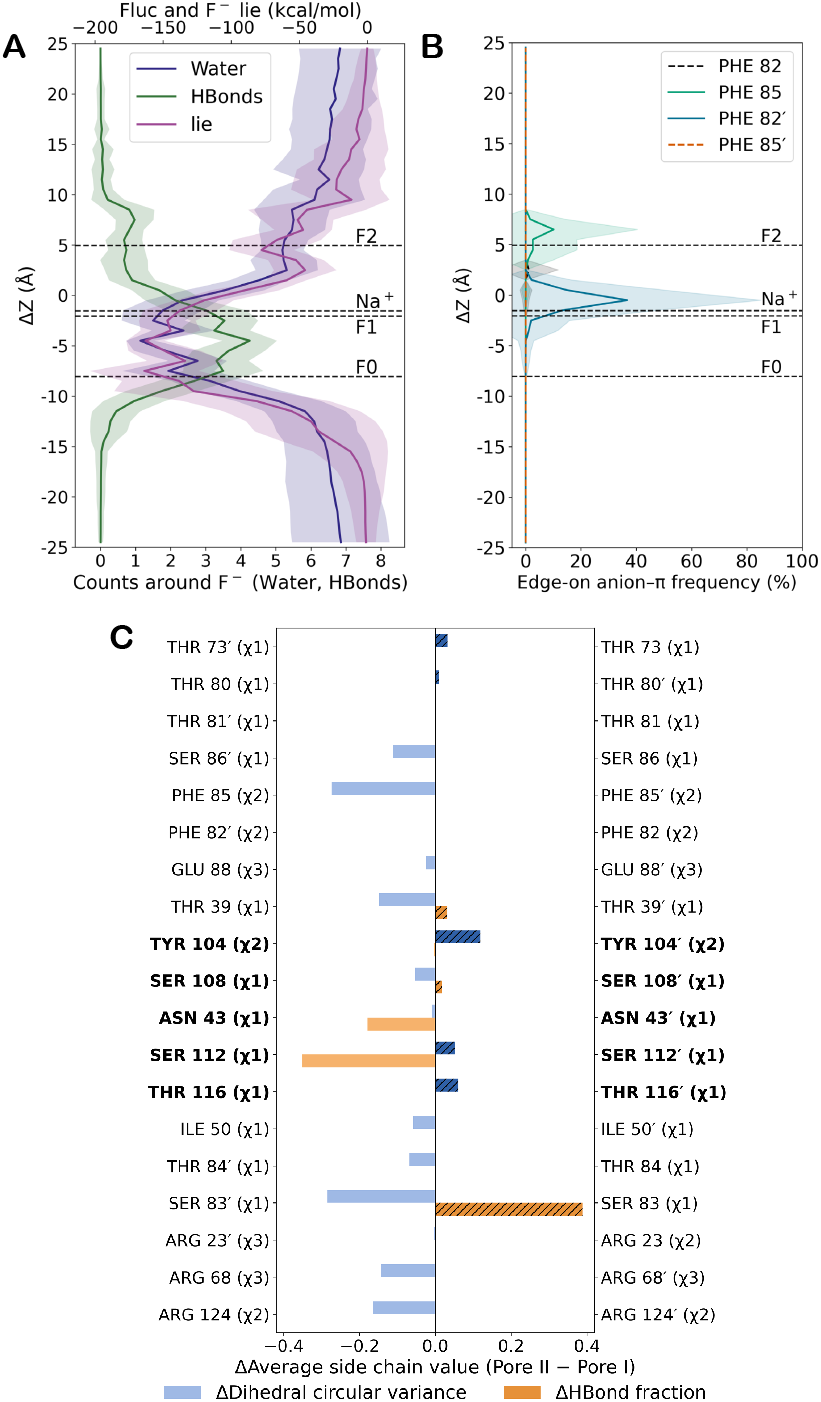
Pore I aligns with the Channsporter mechanism. (*A*) Combined profile of the average number of water molecules within 3.0 Å of fluoride (dark blue), average Fluc–fluoride lie (plum), and average number of hydrogen bonds formed between the protein and fluoride (green) for Pore I–assigned fluoride ions along the bilayer normal (z-axis). (*B*) Frequency of edge-on anion–*π* interactions formed with the phenylalanine residues of the Phe-box for Pore I–assigned fluoride ions along the bilayer normal. Shaded regions in panels A and B indicate the standard deviation of each dataset. Differences in average side-chain circular variance (Δcircular variance; blue) and average fraction of hydrogen bonds with fluoride (ΔHBond fraction; orange) between Pore I-facing residues (left; negative values) and Pore II-facing residues (right; positive values), calculated only for simulations that exhibited fluoride translocation. Polar track residues are bolded for ease of identification.

To further evaluate this mechanistic distinction, we compared Pore I and Pore II in terms of the expected features of a Channsporter mechanism, namely, dynamic side chain rotation coupled to hydrogen-bond-mediated ion conduction. Specifically, we assessed the side chain circular variance (a measure of rotational flexibility) and hydrogen bond fractional occupancy differences between pores for simulations that exhibited fluoride translocation, as reported in Fig. 3C. Most Pore I-facing residues displayed greater side chain mobility than their Pore II counterparts during translocation events (Fig. 3C). However, within the polar track, Pore II-facing residues exhibited slightly higher rotational variance than Pore I-facing residues (Fig. 3C). The average hydrogen bond fractions revealed that the polar track Pore I-facing residues, particularly Asn43 and Ser112, formed more persistent hydrogen bonds with fluoride during translocation than Pore II-facing residues, with remaining residues of the polar track showing similar or marginally higher levels of hydrogen bonding to the fluoride in Pore II (Fig. 3C). The results per pore for these analyses for all simulations and for translocating subsets is provided in Figs. S5–S6.

In summary, the exclusive occurrence of single-ion translocation events in Pore I, combined with its lower overall transport efficiency relative to Pore II, points to a fundamentally different conduction strategy. The evidence supports the Channsporter mechanism, characterized by greater side chain rotational flexibility and longer-lived hydrogen-bonding interactions within the polar track (Fig. 3C), features that likely compensate for the absence of cooperative electrostatic repulsion observed in Pore II.

### Pore II uses the Multi-ion Mechanism for Fluoride Conduction

As described above, fluoride ions were classified into pores based on their predominant location during interactions with Fluc. This approach enabled the identification and analysis of Pore II–assigned fluorides, helping clarify why their behavior differed markedly from Pore I–assigned fluorides.

As with Pore I, the analysis began by examining the hydration number of fluoride ions along the bilayer normal portrayed in Fig. 4A. Owing to the antiparallel orientation of the Fluc homodimer, one might expect an antisymmetric profile relative to Pore I (Fig. 3A). However, the correspondence was not exact. We attribute this deviation to a methodological limitation of the orthogonal-plane classification, which cannot always perfectly distinguish between the lower or intracellular regions of the two pores until a fluoride ion fully commits to one conduction pathway. Nevertheless, the overall trend remained consistent with the expected antisymmetry: fluoride ions dehydrated progressively, reaching a minimum hydration of one to two water molecules near the center of the Pore II upper region (F0^*′*^ and F1^*′*^ containing compartment). As in Pore I, the energy penalty associated with fluoride dehydration (32) was counterbalanced by favorable protein–ion interactions (Fig. 4A). The corresponding hydrogen-bond analysis indicated that this energetic compensation arose primarily from hydrogen bonding, with a maximum of four to five hydrogen bonds formed near the center of the Pore II upper region (Fig. 4A). The first clear divergence between the two pores appeared in the edge-on anion–*π* interactions with the phenylalanine residues of the Phe box as shown in Fig. 4B. In Pore II, these interactions were shallower but more broadly distributed along the conduction pathway. Unlike Pore I, this pattern aligns more closely with the functional pore architecture of the recently determined FEX-*CA* structure (23), in which most polar track residues are replaced by phenylalanines. This correspondence suggests that a broader, more distributed network of edge-on anion–*π* interactions may be functionally advantageous for fluoride conduction in Pore II–like systems. Trajectory inspection revealed that of the nine total fluoride translocation events, six were single-ion permeation events (one per simulation). However, one trajectory, Extracellular 500 mM KF-8e, displayed three translocation events within a single simulation, including two complete (100%) translocations that occurred in rapid succession and in pairs, see Movie S3. These paired translocations provide direct evidence for a Multi-ion conduction mechanism in Pore II.

**Fig. 4.**
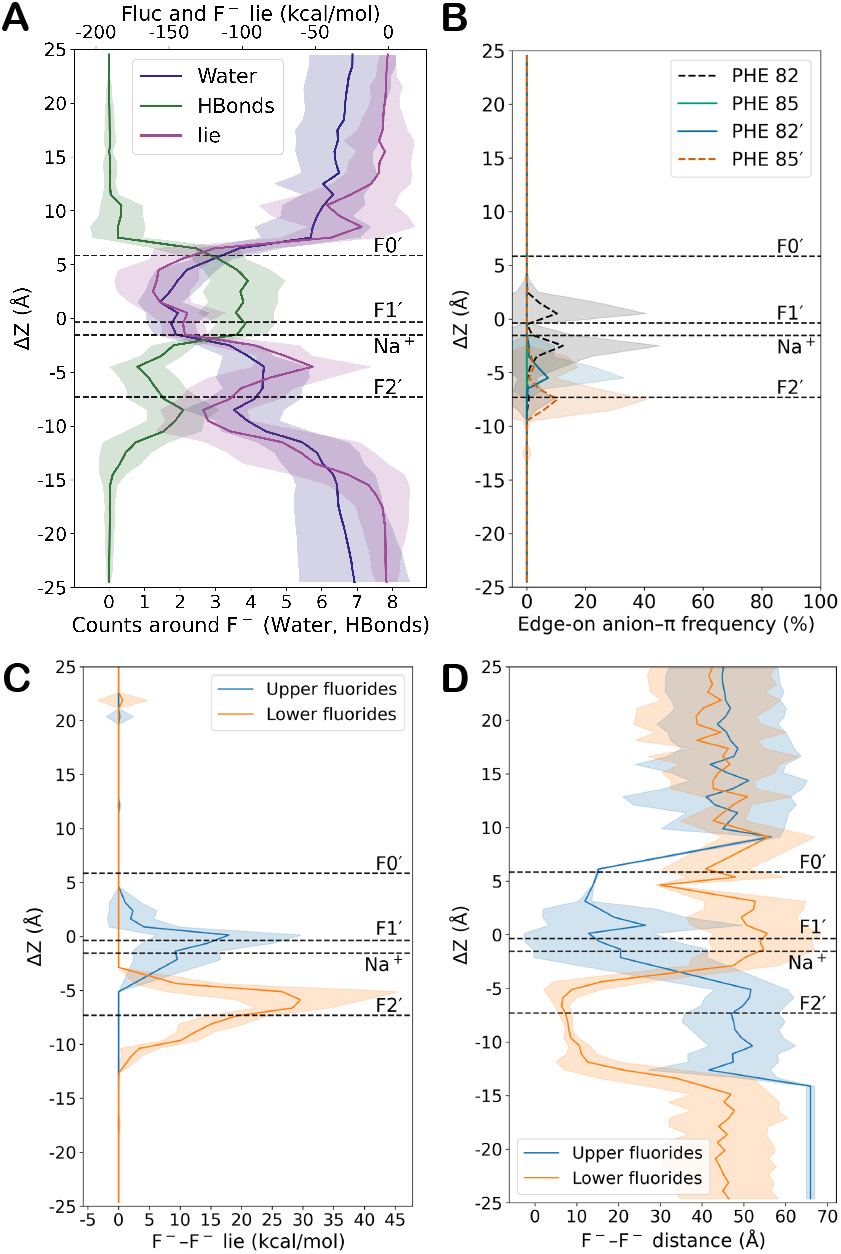
Pore II aligns with the Multi-ion mechanism. (*A*) Combined profile of the average number of water molecules within 3.0 Å of fluoride (dark blue), average Fluc–fluoride lie (plum), and average number of hydrogen bonds formed between the protein and fluoride (green) for Pore II–assigned fluoride ions along the bilayer normal (z-axis). (*B*) Frequency of edge-on anion–*π* interactions formed with the phenylalanine residues of the Phe-box for Pore II–assigned fluoride ions along the bilayer normal. (*C*) Lie profile between the upper and lower fluoride ions (fluoride pair) along the bilayer normal in the simulation that exhibited paired fluoride translocation through Pore II. (*D*) Distance profile between the upper and lower fluoride ions (fluoride pair) along the bilayer normal in the simulation that exhibited paired fluoride translocation through Pore II. Shaded regions indicate the standard deviation of each dataset.

To better characterize this mechanism, we examined the hypothesized electrostatic coupling between cotranslocating fluoride ions, as observed during the rapid paired permeation events in the Extracellular 500mM KF-8e trial. To quantify these interactions, we computed the average lie and distance between the paired fluorides. Fig. 4C shows the average lie profile between the upper and lower fluoride ions along the bilayer normal (z-axis). Interestingly, contrary to the classical alternating-occupancy Multi-ion model, which proposes two sequential electrostatic repulsions (“intracellular F^−^ – F0” and “F0 – F1”) (24), our results revealed a single, well-defined repulsion that aligns with an “F2^*′*^ – *quasi*-F0^*′*^” interaction. This occurred when the upper fluoride in a pair occupied a *quasi*-F0^*′*^ site and the lower fluoride resided at F2^*′*^, as indicated by the high spikes in fluoride-fluoride lie (Fig. 4C). Consistently, the average inter-fluoride distances shown in Fig. 4D display a corresponding single repulsion event, evidenced by a sharp decrease in fluoride-fluoride distance below 10 Å at the same positions. In opposition to the expected symmetry of a single repulsion event, the upper fluoride exhibited lower interaction energies and larger interionic distances at the *quasi*-F0^*′*^ position than the lower fluoride at F2^*′*^. This asymmetry likely reflects the longer residence times of the upper fluoride at the *quasi*-F0^*′*^ site, where it remains poised until a subsequent fluoride arrives from below to provide the electrostatic push required for forward progression. It is noteworthy to mention that the originally proposed alternating mechanism features a structural arrangement with F0 at the intracellular side and F2 at the extracellular side (24, 25). This configuration corresponds to the orientation of Pore I in our model, rather than to the F2^*′*^ intracellular / F0^*′*^ extracellular arrangement observed for Pore II. More broadly, these findings reinforce that, owing to their antisymmetry, the two pores likely do not operate under a single, unified conduction strategy; and that applying one mechanistic framework to both may overlook fundamental functional distinctions. Together, these observations support a mechanism in which a single, dominant electrostatic repulsion event drives conduction in Pore II, consistent with a Multi-ion conduction model.

In summary, the occurrence of paired electrostatic repulsion events in Pore II, combined with its higher overall transport efficiency relative to Pore I, indicates a distinct conduction strategy. The evidence supports the Multi-ion mechanism, characterized by electrostatic repulsion between sequential fluoride ions that reduces the reliance on long-lived hydrogen-bond networks and coordinated side chain rotations observed in the Channsporter-type Pore I.

### Role of the Central Sodium Ion in Fluoride Transport

During trajectory inspection, we observed two rare events in which the central sodium ion escaped from the Fluc protein. Fig. S7A–C show the typical secondary structure surrounding the sodium-binding site, closely resembling the crystal structures. In contrast, two simulations exhibited clear sodium release from this site as displayed in Fig. S7D–F. This observation was unexpected, as previous experimental work by McIlwain *et al*. demonstrated that sodium removal from this position is experimentally difficult (21). In one of the sodium-escape trajectories, a pronounced distortion of the local secondary structure was observed, which likely weakened the backbone coordination of the sodium ion and facilitated its release (Fig. S7D–E). Specifically, while the loop coordinating sodium in Chain A retained a fold similar to the crystal structure, Chain B exhibited substantial unfolding in the proximal segment of the transmembrane helix 3b^*′*^ to the loop, disrupting the backbone–sodium coordination at the helix break due to increased loop flexibility (Fig. S7E). Interestingly, in the other escape trajectory, no significant secondary structure changes were detected near the binding site, with the exception of the region surrounding F74^*′*^ (helix 3a^*′*^), which became more structured and rigid (Fig. S7F). This observation leaves the open question of what structural or dynamic features are truly critical for sodium retention. These escape events prompted a more detailed investigation of sodium ion dynamics. As attested by Fig. 2C, the central sodium exhibits a surprisingly large range of motion, sampling a roughly spherical volume with a radius of 2.5 Å. Such extensive mobility likely explains why the ion is stabilized by a relatively loose coordination environment provided by the four protein backbone carbonyls complemented by water molecules as seen in Fig. S8 and consistent with the total coordination number of 5–6 reported by Yue *et al*. in Fluc-*Ec2* and Kang *et al*. in FEX-*CA*(23, 27). To understand the functional implications of this mobility, we closely examined the trajectories to characterize sodium behavior during transport.

In Pore I, a recurring sequence of sodium movements was identified and shown in Fig. 5A. Initially, the sodium ion shifted downward toward the intracellular side, appearing to assist in the recruitment of fluoride from the intracellular medium. It then returned toward the central position, guiding the fluoride upward along the conduction pathway. Finally, the sodium moved further toward the extracellular side, continuing to escort the fluoride toward the channel exit. To better characterize the sodium positions associated with fluoride translocation, we mapped the sodium coordinates sampled during Pore I translocation events highlighted in cyan in Fig. S9A. These highlighted positions aligned closely with the average lie profile between sodium and Pore I-assigned fluorides in Fig. 5B, which reveals a strong interaction centered in the Pore I lower region corresponding to the *quasi*-F0 site. Besides, sodium also samples the upper range of its accessible volume, where its distance from fluoride increases, likely weakening the cation–anion association and facilitating fluoride release toward the extracellular side. These findings suggest that sodium primarily acts as a recruiting agent in Pore I, stabilizing the entry of fluoride ions. However, this high-affinity interaction likely hinders subsequent progression of the ion along the pore, providing a possible explanation for the lower transport efficiency of Pore I compared to Pore II. As in other membrane transporters, effective function requires balancing substrate affinity with rapid turnover. In Pore I, excessive binding strength at the entry site may compromise overall conduction speed.

**Fig. 5.**
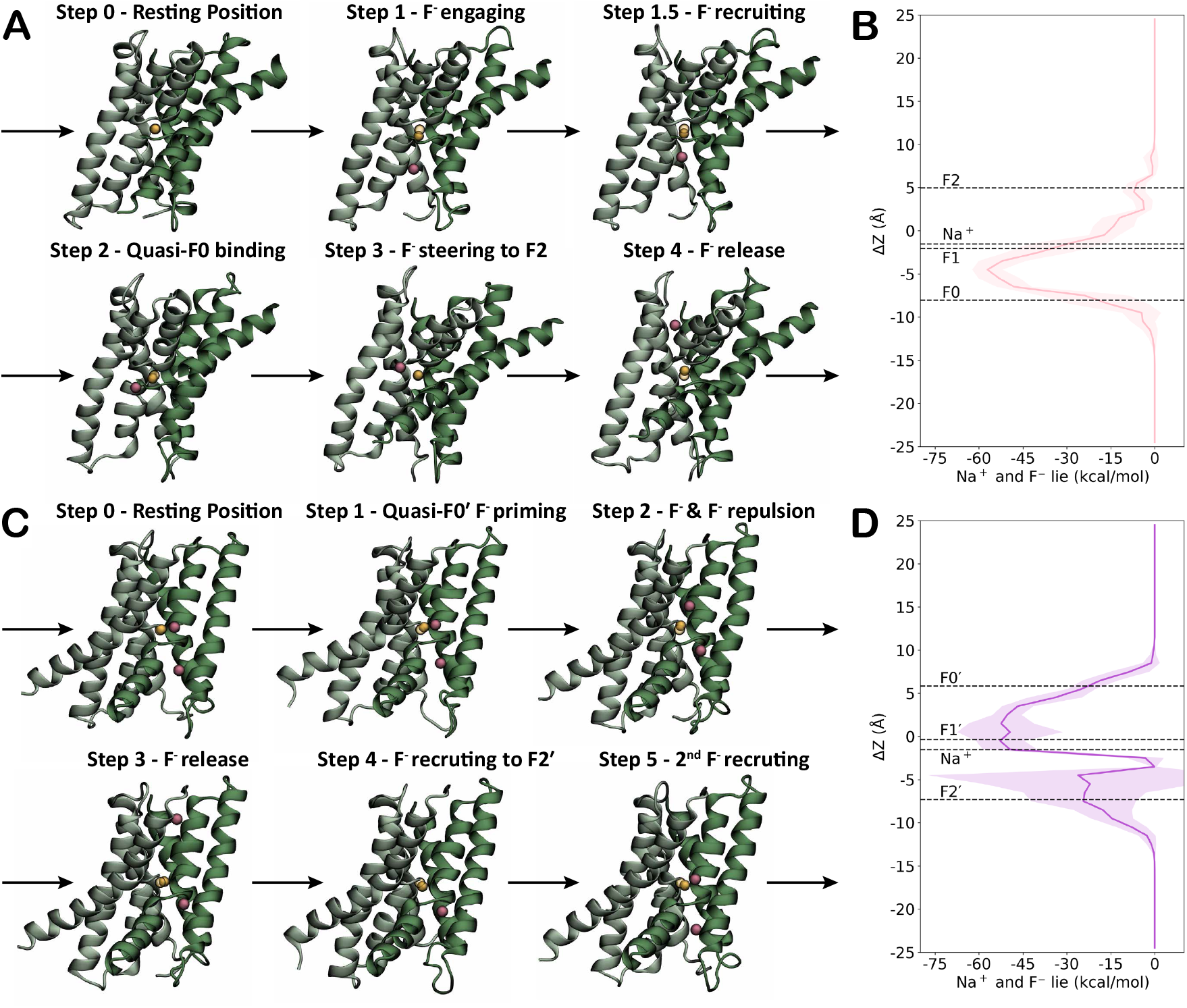
Central sodium movement is pivotal for fluoride conductivity in Fluc. (*A*) Stepwise schematic of the proposed role of the central sodium ion during Pore I fluoride transport, based on representative simulation snapshots. *B*) Lie profile between the central sodium ion and Pore I-assigned fluoride ions along the bilayer normal (z-axis); shaded regions denote the standard deviation. (*C*) Stepwise schematic of the proposed role of the central sodium ion during Pore II fluoride transport, based on representative simulation snapshots. (*D*) Lie profile between the central sodium ion and Pore II-assigned fluoride ions along the bilayer normal, with shaded regions indicating the standard deviation. Transmembrane helices 3b and 3b^*′*^ have been omitted accordingly from the graphical representation for ease of visualization of the ions.

Similarly, sodium involvement in Pore II was characterized, with particular attention to the trajectory that featured three translocation events (two of which were 100%). As shown in Fig. 5C, a recurring pattern of sodium–fluoride interactions also emerged in this pore. Initially, an upper fluoride ion engaged strongly with the central sodium at a quasi-F0^*′*^ position. This interaction persisted until a second fluoride approached from below, whose electrostatic repulsion displaced the upper ion toward the extracellular media. Release from this site appeared to require two cooperative factors: continued repulsion from the following fluoride and a downward movement of sodium, which disengaged from the upper fluoride and reset the system to recruit the next incoming ion. Thus, the sodium exhibited a downward movement to recruit an incoming fluoride to subsequently shift upward to escort the fluoride toward the electrostatic priming *quasi*-F0^*′*^ site to restart the cycle. The sodium positions associated with Pore II translocation events mapped in Fig. S9B revealed a spatial pattern remarkably similar to that observed for Pore I (Fig. S9A). These findings imply that, under physiological conditions, the two pores may be capable of translocating ions concurrently rather than sequentially, with neither pore dependent on the completion of the other’s conduction cycle. As in Pore I, the sodium positions associated with translocation align closely with the sodium and Pore II-assigned fluorides average lie profile displayed in Fig. 5D. However, in contrast to Pore I, Pore II displays two well-defined interaction sites: a weaker intracellular interaction at F2^*′*^, which facilitates initial fluoride recruitment, and a stronger interaction within the upper region at the *quasi*-F0^*′*^ site, which guides fluoride toward the extracellular side. Because this upper site exhibits a strong interaction, fluoride can become transiently trapped, necessitating electrostatic repulsion from a trailing ion to complete translocation, consistent with the Multi-ion model.

Together, these results reveal contrasting roles of sodium in the two pores. In Pore I, the strongest sodium–fluoride interaction occurs in the intracellular entrance (Pore I lower region), where sodium tightly stabilizes the incoming fluoride. However, because this interaction site lies far from the extracellular exit, fluoride must traverse nearly the entire length of the pore prior to release, thereby slowing and reducing the efficiency of transport. In contrast, in Pore II, the strongest sodium–fluoride interactions occur near the extracellular side (Pore II upper region). Here, sodium acts as a dynamic cofactor that positions fluoride in place for electrostatic repulsion by a trailing ion, enabling more efficient forward progression toward the extracellular medium.

## Discussion

The fluoride channel Fluc is an idiosyncratic and remarkable protein, yet several aspects of its mechanism remain unresolved. How it achieves exceptionally high selectivity despite conduction rates comparable to the fastest ion channels has been a long-standing question. Likewise, the precise mechanism of fluoride translocation and the functional role of the central sodium ion have remained elusive. In this theoretical study, we used long-timescale CompEL simulations to address these questions, offering mechanistic insights that can guide future experimental validation.

We propose that Fluc does not operate via a single mechanism, but instead employs two distinct yet complementary modes of action, one in each pore, arising from the antiparallel, structurally antisymmetric assembly of the homodimer. Our results show that Pore I conforms more closely to the Channsporter mechanism originally proposed by Stockbridge *et al*. (17). During the simulations, this pore exhibited slower, single-ion translocation events and longer F^−^ residence times near the intracellular entrance, where the strongest sodium–fluoride interactions occurred. The strong attraction at the entrance of the pore before traversing nearly the entire pore likely reduces overall transport efficiency. In contrast, Pore II behaves according to the Multi-ion conduction mechanism, consistent with experimental observations by McIlwain *et al*. (24) and Zhang *et al*. (25). In this pore, the strongest sodium–fluoride interactions occurred near the extracellular side, where paired fluoride ions cooperated through electrostatic repulsion, facilitating rapid and sequential release. This mechanistic and positional asymmetry provides a compelling explanation for the differences in efficiency between the two pores observed during the simulations. Moreover, it offers an evolutionary rationale for why the eukaryotic Fluc homolog, FEX, has apparently deprecated one pore, possibly optimizing the faster, Multiion–type pathway represented by Pore II. Experimental evidence from Kang and coworkers (23) confirms that FEX adopts an intracellular-to-extracellular bilayer orientation that aligns Fluc’s less efficient Pore I with the vestigial pore, whereas the active pore corresponds to Fluc’s more efficient Pore 2. This alignment supports the hypothesis that Pore I was evolutionarily silenced while Pore II was retained and refined for transport. Whether this optimization primarily enhanced transport rate or anion selectivity remains an open question.

Our findings also suggest that the central sodium ion plays an active functional role in fluoride transport. The sodium ion exhibits a substantial range of motion within its binding site and engages dynamically with fluoride ions during translocation. In Pore I, sodium acts primarily as a high-affinity recruiter, guiding fluoride ions from the intracellular side but at the expense of slower release. In Pore II, sodium behaves as a priming or holding cofactor that holds the leading fluoride in place for efficient fluoride–fluoride electrostatic repulsion and propels the leading ion outward. This dynamic role provides a possible explanation for experimental observations that lithium substitution inactivates Fluc: lithium’s preference for four-fold coordination likely leads to a tighter, less flexible binding mode, impairing its ability to participate in the transport cycle (21, 23). Although we did not observe evidence clarifying sodium’s potential role in structural stability or dimer assembly, our results support the opposite interpretation, namely, that Fluc’s architecture has evolved around sodium’s mechanistic contribution to transport, not vice versa, just as the work of Ernst *et al*. suggests (33).

With respect to fluoride selectivity, our results show that the ion becomes almost completely dehydrated within the high-affinity regions of both pores. However, dehydration alone does not fully explain the extraordinary selectivity of Fluc. Given the observed coupling between sodium and fluoride, we hypothesize that changes in the identity or coordination of the central cation could modulate substrate selectivity, potentially enabling transport of alternative anions. To our knowledge, this possibility remains experimentally untested. While previous studies have examined the effect of substituting different cations at the central site on fluoride conductivity (21, 23), they did not assess whether such substitutions permit the conductivity of other anions or substrates.

In conclusion, our work provides a unified model of asymmetric dual-pore conduction in Fluc, integrating long-timescale electrophysiological simulations with mechanistic insight. The results highlight two divergent transport strategies operating within a single homodimeric framework and identify sodium as a central, mechanistically active cofactor in fluoride translocation. We anticipate that forthcoming experimental validation will further refine this model and open new avenues in membrane protein engineering, where dual-pore architectures may be exploited for selective or coupled transport of different substrates.

## Materials and Methods

### Dual Bilayer System Preparation

Considering that the Fluc protein is an exceptionally fast ion channel, conducting approximately one translocation per microsecond, or 10^6^ translocations per second (10, 26, 31), we determined that long-timescale equilibrium Computational electrophysiology (CompEL) simulations (29, 30) would be the most appropriate approach to study this system in an unbiased manner. For this purpose, the high-resolution crystal structure of *Bordetella pertussis* (Bpe) Fluc (PDB ID: 5NKQ) (17), which features a centrally resolved sodium ion (Na^+^) and fluoride ions (*F*^−^) occupying key binding sites was selected. The system preparation protocol was as follows. First, protonation states for all titratable residues were assigned at pH 7.4 using the H++ server (34–36). Both monomers were capped using an acetyl group at the N-termini and an N-methyl group at the C-termini using the tleap module from the Amber20 suite (37). Subsequently, the system was embedded in a POPC lipid bilayer and solvated with 0.15 M KCl using packmol-memgen from the Amber20 suite (37–39). KCl was chosen instead of NaCl to avoid competition with the centrally bound Na^+^ ion. To simulate an electrochemical potential across the membrane, we constructed a second bilayer system lacking the protein also using packmol-memgen, and positioned it below the protein-containing bilayer to form a dual-membrane setup suitable for electrophysiological conditions. To further assess how different electrochemical gradients might influence fluoride transport, we simulated each system under three intracellular KF concentrations: 0.15 M, 0.25 M, and 0.5 M, and three corresponding charge imbalances (0e, 4e, and 8e), achieved by displacing cations from the intracellular compartment to the extracellular space for each concentration. Fig. S1A and Table S1 present a graphical representation of the dual bilayer system and a summary of the different electrochemical gradient configurations. Further details on the construction of the double-bilayer system and related analyses are provided in this book chapter (40). To enhance sampling and analyze the ongoing debate regarding Fluc’s fluoride crystallized locations (22, 25), six distinct initial fluoride configurations were prepared: (1) fully occupied, (2) empty, one pore occupied, (4) intracellular sites occupied, (5) extracellular sites occupied, and (6) diagonally occupied, where one F^−^ resides at the intracellular site of one channel and another at the extracellular site of the other. See Fig. S1B for a visual summary of these initial states. This setup yielded a total of 54 distinct simulation trials. Each system, with its respective fluoride configuration and KF concentration, was processed through tleap to generate parameter and coordinate files. The simulations employed the ff14SB force field for proteins, lipid17 for lipids, and TIP3P for water molecules (41–44), which also provides parameters for the counterions and for fluoride.

### Computational Electrophysiology (CompEL) Simulations

The dualmembrane systems were subjected to molecular dynamics (MD) simulations using the Amber20 suite (37). Initially, energy minimization was performed in three consecutive stages of 10,000 steps each. For each stage, the first 5,000 steps used the steepest descent algorithm, followed by 5,000 steps of the conjugate gradient algorithm. The minimization protocol involved a gradual reduction of positional restraints: in the first stage, all atoms except water and hydrogen atoms were restrained with a force constant of 7.5 kcal/mol/Å^2^; in the second stage, only the protein atoms were restrained with the same force constant; and in the final minimization, all restraints were removed. Following minimization, the systems were heated from 0 K to 100 K and then from 100 K to 303 K in two successive 10 ps intervals under the NVT ensemble. During heating, all protein and lipid atoms were restrained with a force constant of 5.0 kcal/mol/Å^2^. Subsequently, the systems underwent a membrane equilibration under the NPT ensemble, consisting of 10 cycles of 500 ps each. This step was essential for stabilizing the periodic boundary conditions (PBCs) and ensuring appropriate membrane packing. Finally, production runs were carried out for each system for a total of 2 *µ*s under the NPT ensemble. Simulations were conducted using a 2 fs time step, Langevin dynamics for temperature control, Berendsen barostat for pressure control, the SHAKE algorithm to constrain bonds involving hydrogen atoms, and a nonbonded interaction cutoff of 12 Å (45, 46). All simulations were executed using the GPU-accelerated *pmemd* engine in the Amber20 suite (37).

### MD Trajectory Analysis

The total simulation time amounted to 108 *µ*s, comprising 54 distinct system configurations, each simulated for 2 *µ*s. The first 100 ns of every trajectory were excluded from analysis to allow for equilibration, as confirmed by the convergence of the backbone RMSD over time (Fig. S2A). Trajectory analyses were performed using a combination of established and in-house tools. Most analyses were conducted with CPPTRAJ, part of the AmberTools package in the Amber20 suite (37, 47, 48), including calculations of RMSD, RMSF, salt bridges, hydrogen bonds, side chain dihedral angles, secondary structure, water hydration, interatomic distances, atom locations, and Linear Interaction Energy (lie). Additional custom analyses were carried out using Tcl and Python scripts developed in-house and tailored to the specific dynamic and structural features of the Fluc system. Tcl and Python scripting was used to quantify fluoride ion positions via a set of orthogonal planes, enabling pore assignment and regional classification. Python scripts were used for post-processing tasks such as computing pore occupancy and translocation frequencies, quantifying edge-on anion–*π* interactions, and generating all graphical representations. Molecular visualizations and structure inspections were performed using VMD (49), with rendering via the Tachyon ray tracer (50).

The edge-on anion–*π* interaction analysis followed the definition by Yue *et al* (27). Two geometric criteria were required for an interaction to be counted: (1) the distance between the fluoride ion and any phenylalanine ring carbon was less than 4.5 Å, and (2) the angle between the aromatic ring plane and the fluoride position, derived from the angle between the ring normal and the vector connecting the ring center to the fluoride, was less than 35◦.

For Fig. 2C, and Fig. S9 three-dimensional Cartesian coordinates (*x, y*) were reduced to a single projected coordinate to generate two-dimensional views, using the transformation *u* = *x* cos *ϕ* + *y* sin *ϕ*, where *ϕ* (in degrees) was selected to reproduce the perspective of the 3D visualizations.

## Data, Materials, and Software Availability

The MD simulation input files, one representative trajectory for each conducting mechanism, and the inhouse data-processing and plotting codes have been uploaded to Zenodo: 10.5281/zenodo.17982196. These files can be accessed through the following link: https://tinyurl.com/ef5uc77n.

## ACKNOWLEDGMENTS

The authors thank the Office of Information Technology and the Cyber Infrastructure Research Computing (CIRC) at the University of Texas at Dallas, as well as the Texas Advanced Computing Center (TACC) at the University of Texas at Austin, for providing HPC resources that have contributed to the research results reported within this paper.

The research reported in this publication was supported by the National Institutes of Health under award number R35GM155106 (to H.T.) and the University of Texas at Dallas. The content is solely the responsibility of the authors and does not necessarily represent the official views of the National Institutes of Health.

