## Supplemental Information for "Asymmetry-induced distinct mechanisms and the transporting role of sodium in bacterial fluoride channel Fluc"

### This PDF file includes:

Figs. S1 to S9
Tables S1 to S2
Legends for Movies S1 to S3

### Other supporting materials for this manuscript include the following:

Movies S1 to S3

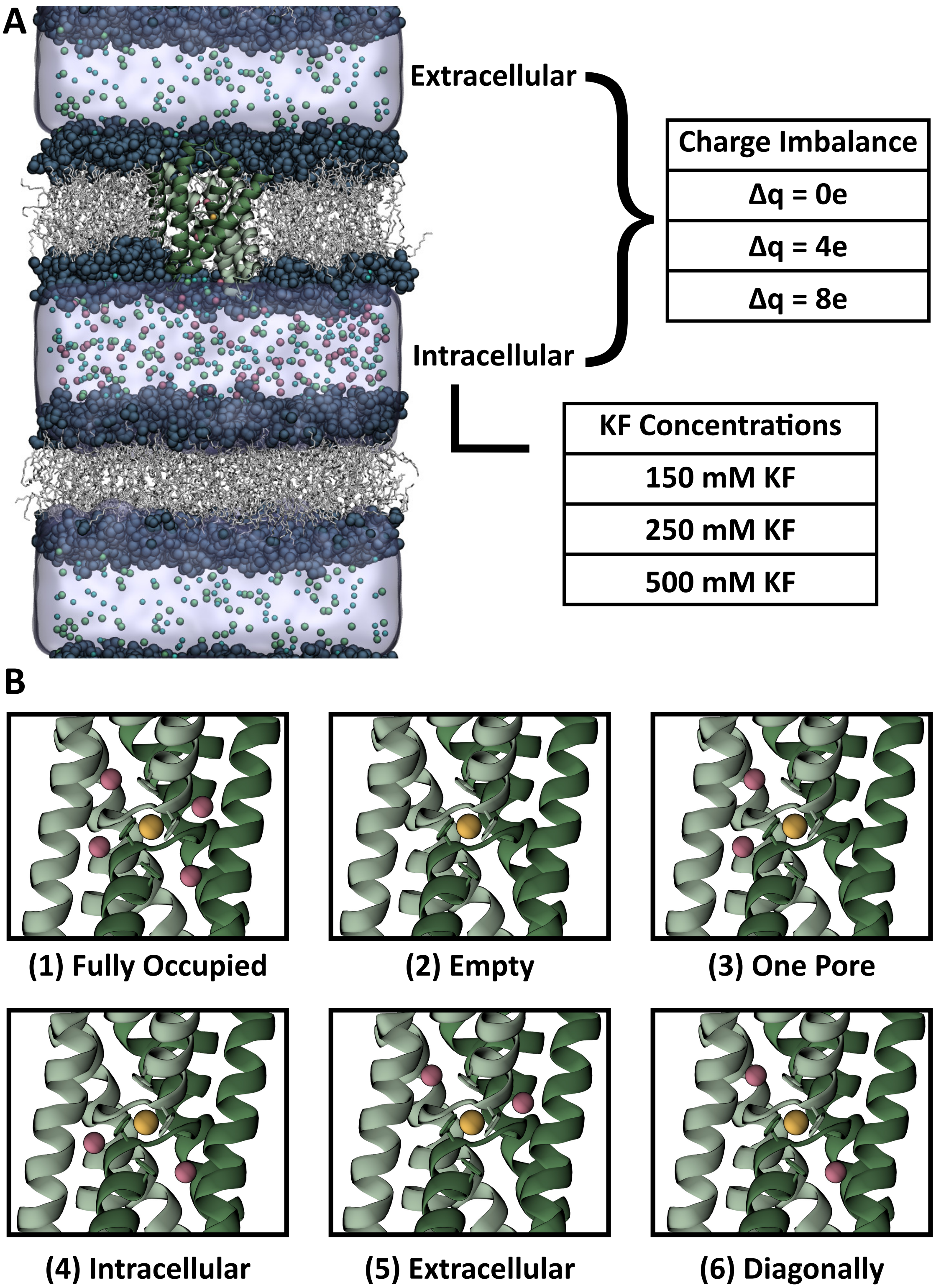


**Fig. S1.** CompEL setup. (*A*) Dual-bilayer system used for MD production with the Fluc protein embedded in one of the bilayers. On the right, tables summarize the combinations of charge imbalances (0e, 4e, 8e) and intracellular KF concentrations (0.15 M, 0.25 M, and 0.5 M) used to establish different electrochemical gradients and enhance sampling. (*B*) The six initial fluoride-binding configurations tested: (1) fully occupied, (2) empty, (3) one pore occupied, (4) intracellular sites occupied, (5) extracellular sites occupied, and (6) diagonally occupied (one $F^-$ in the intracellular site of one pore, the other in the extracellular site of the second pore). The Fluc homodimer is shown in two shades of green. Lipid acyl chains are white, polar headgroups are dark blue, and ions are represented as spheres: sodium (yellow), fluoride (pink), potassium (light blue), and chloride (green).

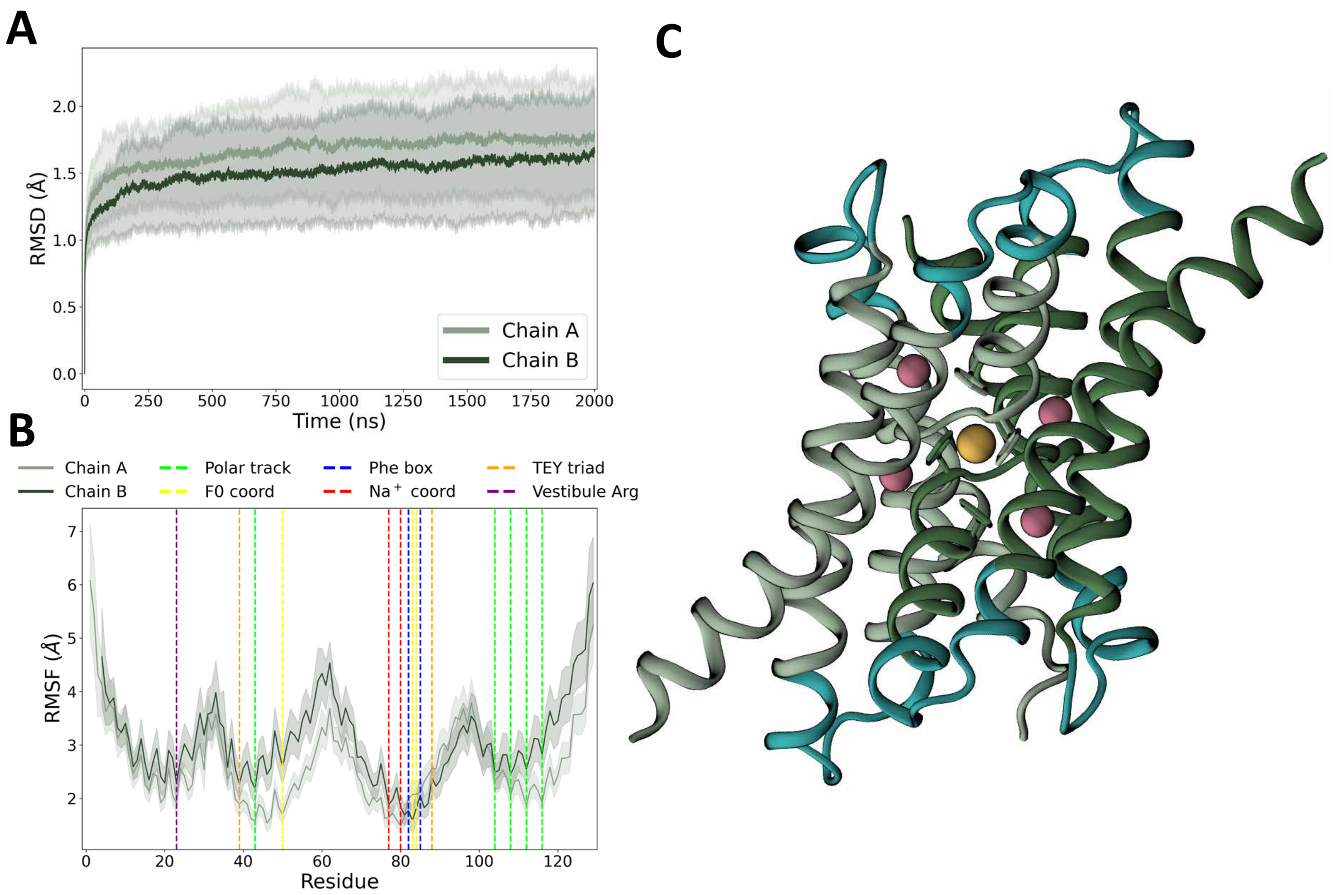


**Fig. S2.** RMS Analysis of Fluc homodimer for all systems. (*A*) Average backbone RMSD relative to the initial frame for each trial, calculated separately for each monomer. Standard deviation is represented as the shaded areas around the curves of each chain. (*B*) Average per-residue all-atom RMSF values, also analyzed per monomer, with annotations highlighting key residues for fluoride conductivity and sodium coordination. As before, the standard deviation is the shaded area around each curve. (*C*) Fluc dimer structure with cyan highlights indicating highly dynamic regions, corresponding to residues with elevated RMSF values observed across trials.

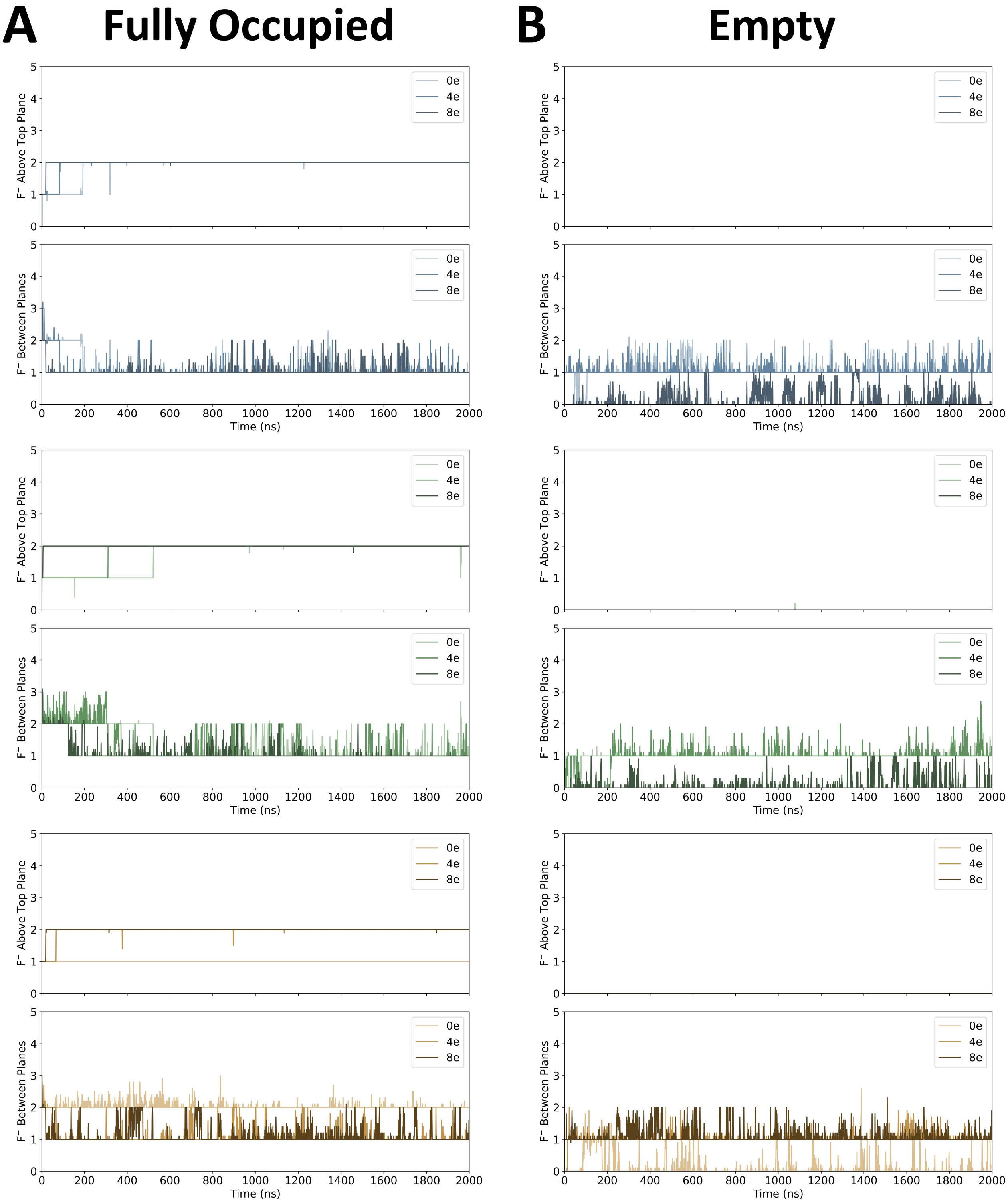
A
Fully Occupied
B
Empty
F⁻ Above Top Plane
F⁻ Between Planes
0e
4e
8e
Time (ns)

**C** **One Pore** **D** **Intracellular**

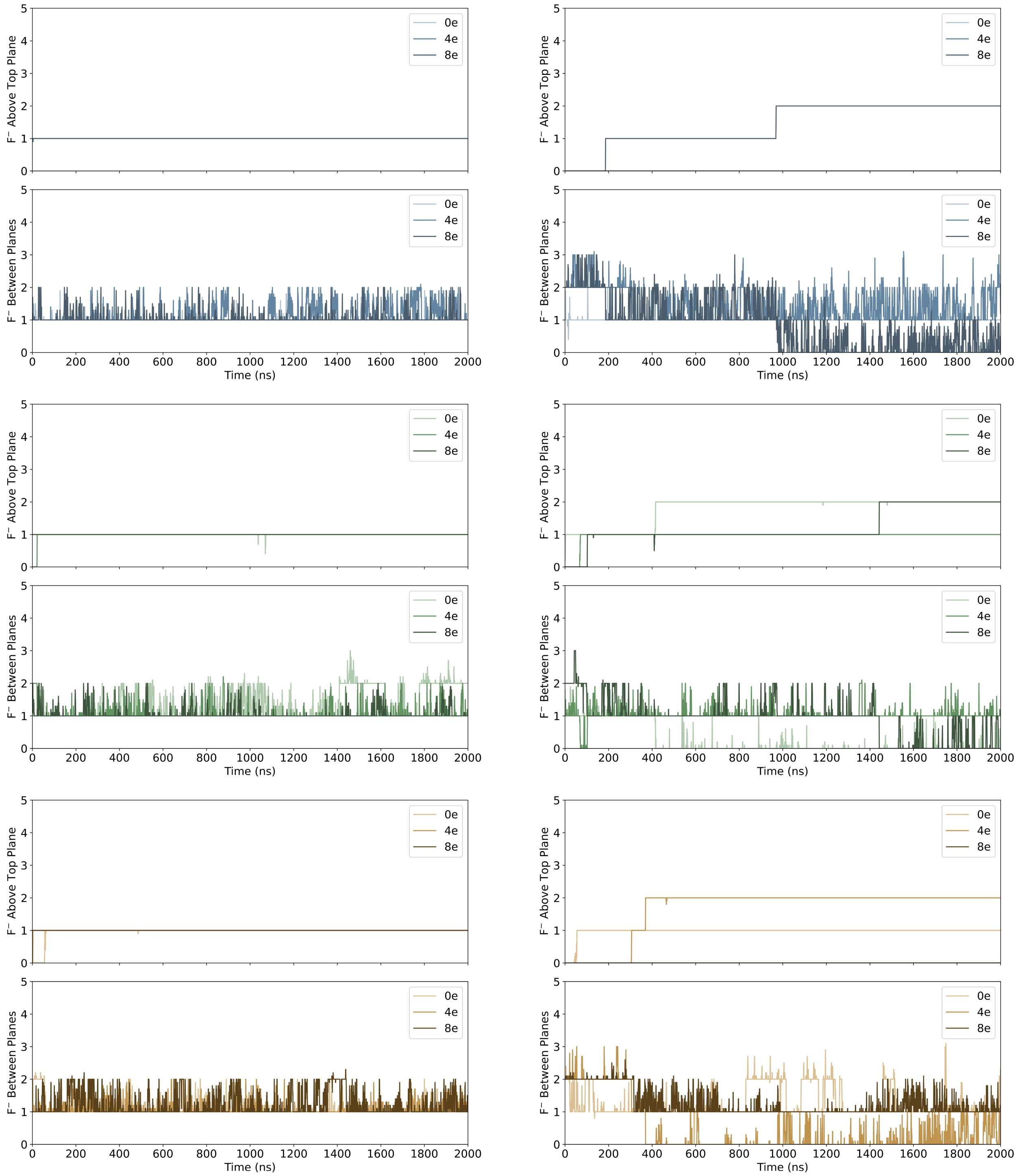

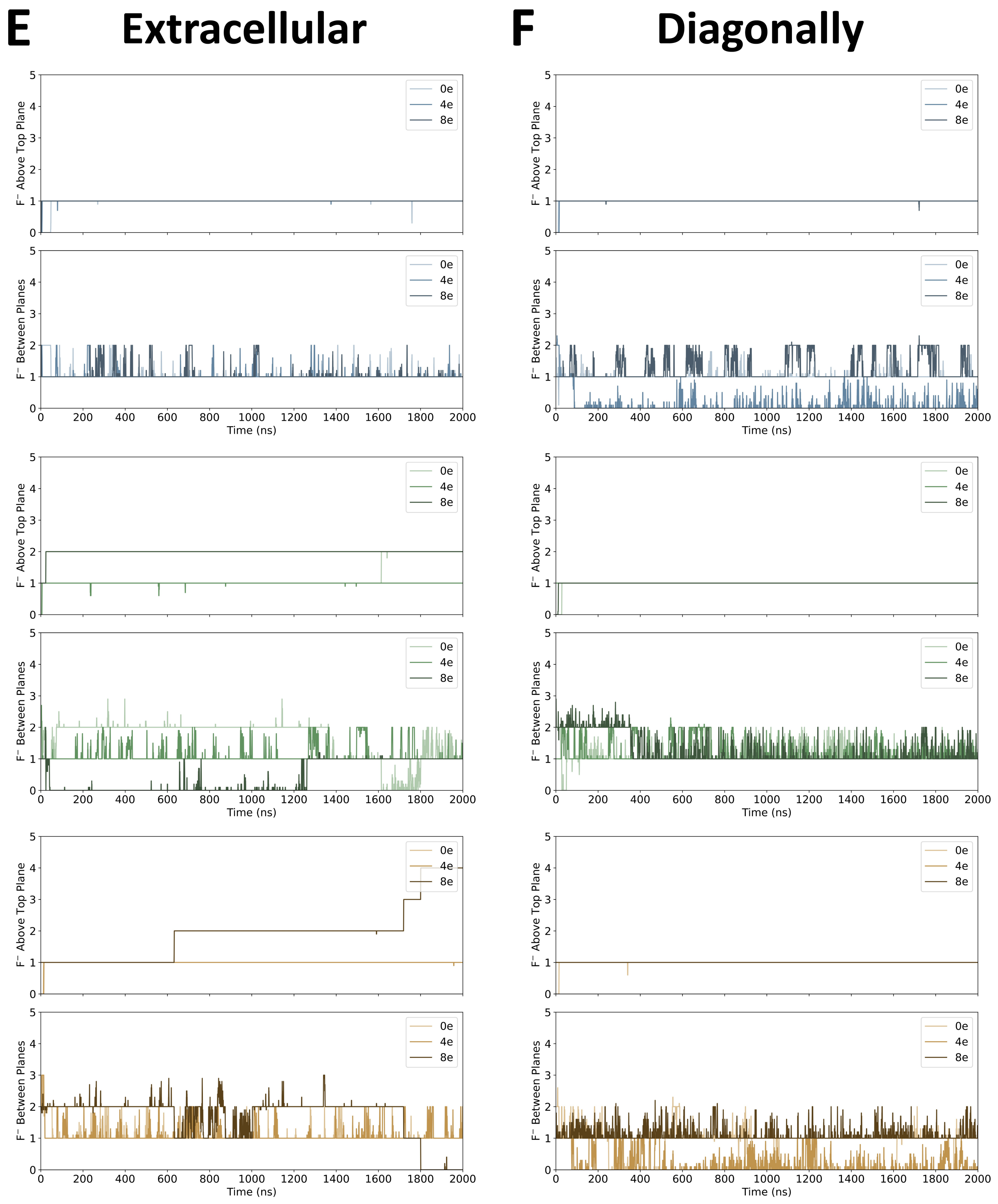


**Fig. S3.** Fluoride pore occupancy and translocation quantification. Results of the in-house Tcl script that employed orthogonal planes to analyze fluoride content within the Fluc structure and at the extracellular media for (*A*) fully occupied, (*B*) empty, (*C*) one pore occupied, (*D*) intracellular sites occupied, (*E*) extracellular sites occupied, and (*F*) diagonally occupied. The color trend from blue to green to brown represents the three different KF concentrations used, 0.15 M, 0.25 M, and 0.5 M, respectively. The charge imbalances are the different curves represented by the different color tones of their respective KF concentration, with lighter tones representing lower charge imbalances and darker tones of the color larger charge imbalances. To enhance the readability of the plots, the data were smoothed using a running average.

 

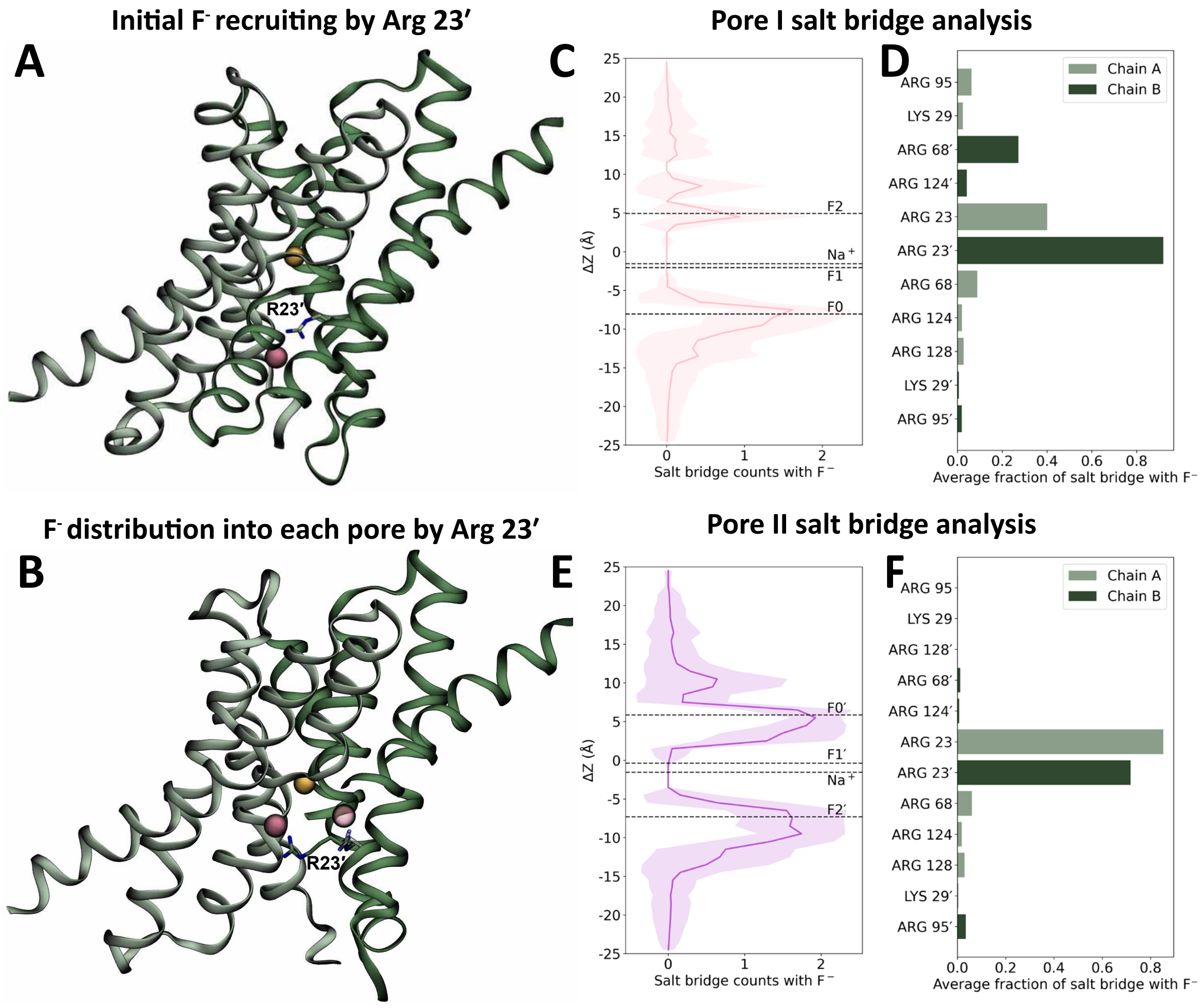


**Fig. S4.** Initial recruitment and distribution of intracellular fluoride ions by Arg23′ in Fluc. (*A*) Electrostatic attraction between the positively charged guanidinium group of Arg23′ and the negatively charged intracellular fluoride initiates the recruitment process. (*B*) Subsequent rotation of the Arg23′ side chain enables fluoride distribution into both pores. For visual clarity, two instances of this event are shown simultaneously: fluoride distribution into Pore I is rendered fully opaque, whereas Pore II is shown transparently. Transmembrane helices 3b′ and 4′ has been omitted accordingly from the graphical representation for ease of visualization of the ions. (*C*) Average number of salt bridges between fluoride and Fluc residues along the bilayer normal (z-axis) for Pore I across all simulations; shaded regions represent the standard deviation. (*D*) Average fraction of individual salt bridges formed between fluoride and Fluc residues in Pore I. (*E*) Average number of fluoride–Fluc salt bridges along the bilayer normal for Pore II, with standard deviations shown as shaded regions. (*F*) Average fraction of fluoride–Fluc salt bridges per residue for Pore II.

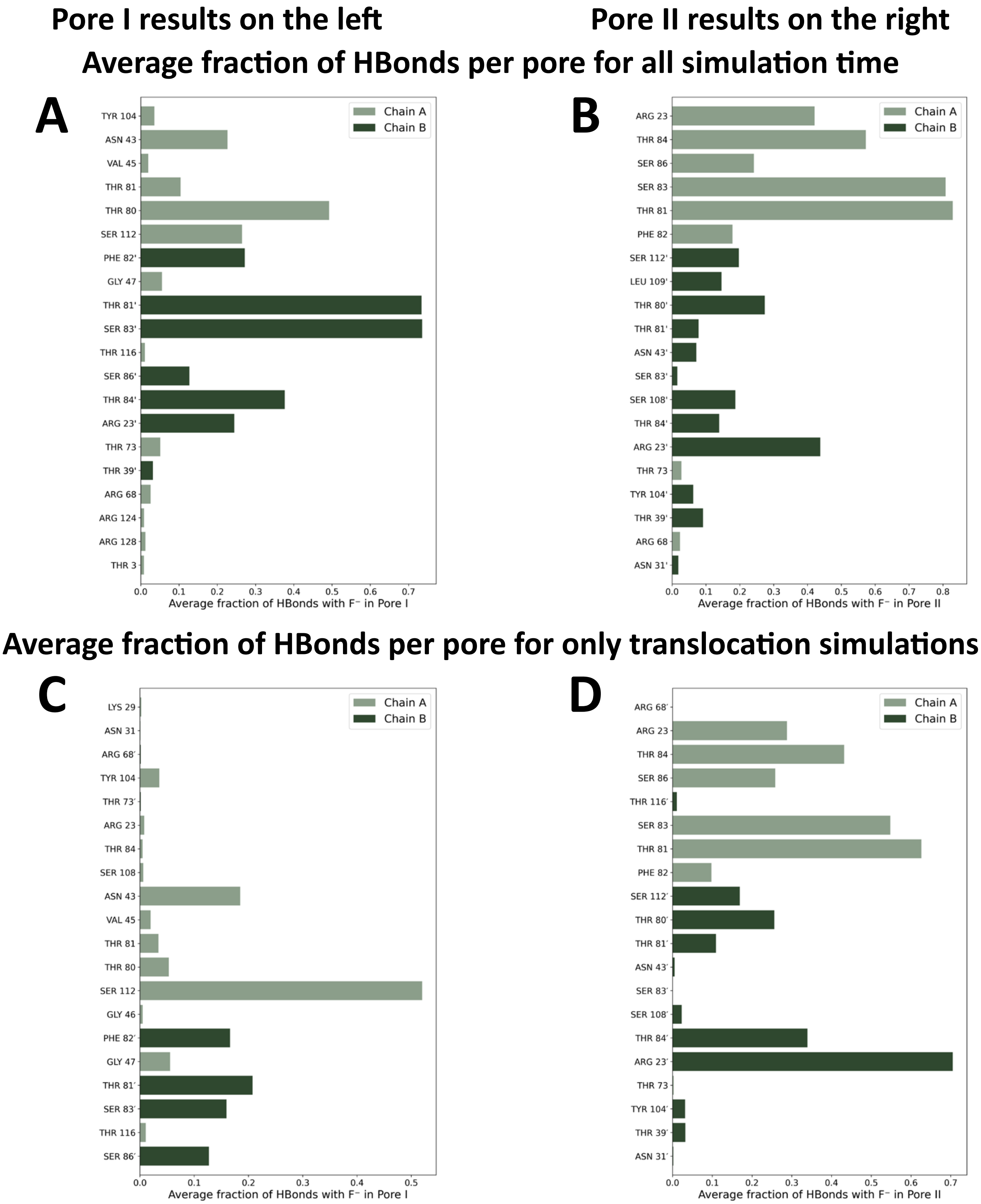


**Fig. S5.** Average fraction of hydrogen bonds per pore. (*A*) Average fraction of hydrogen bonds between fluoride ions and Fluc residues in Pore I over the entire simulation time. (*B*) Average fraction of hydrogen bonds between fluoride ions and Fluc residues in Pore II over the entire simulation time. (*C*) Average fraction of hydrogen bonds for Pore I calculated only from simulations that exhibited fluoride translocation. (*D*) Average fraction of hydrogen bonds for Pore II calculated only from simulations that exhibited fluoride translocation.

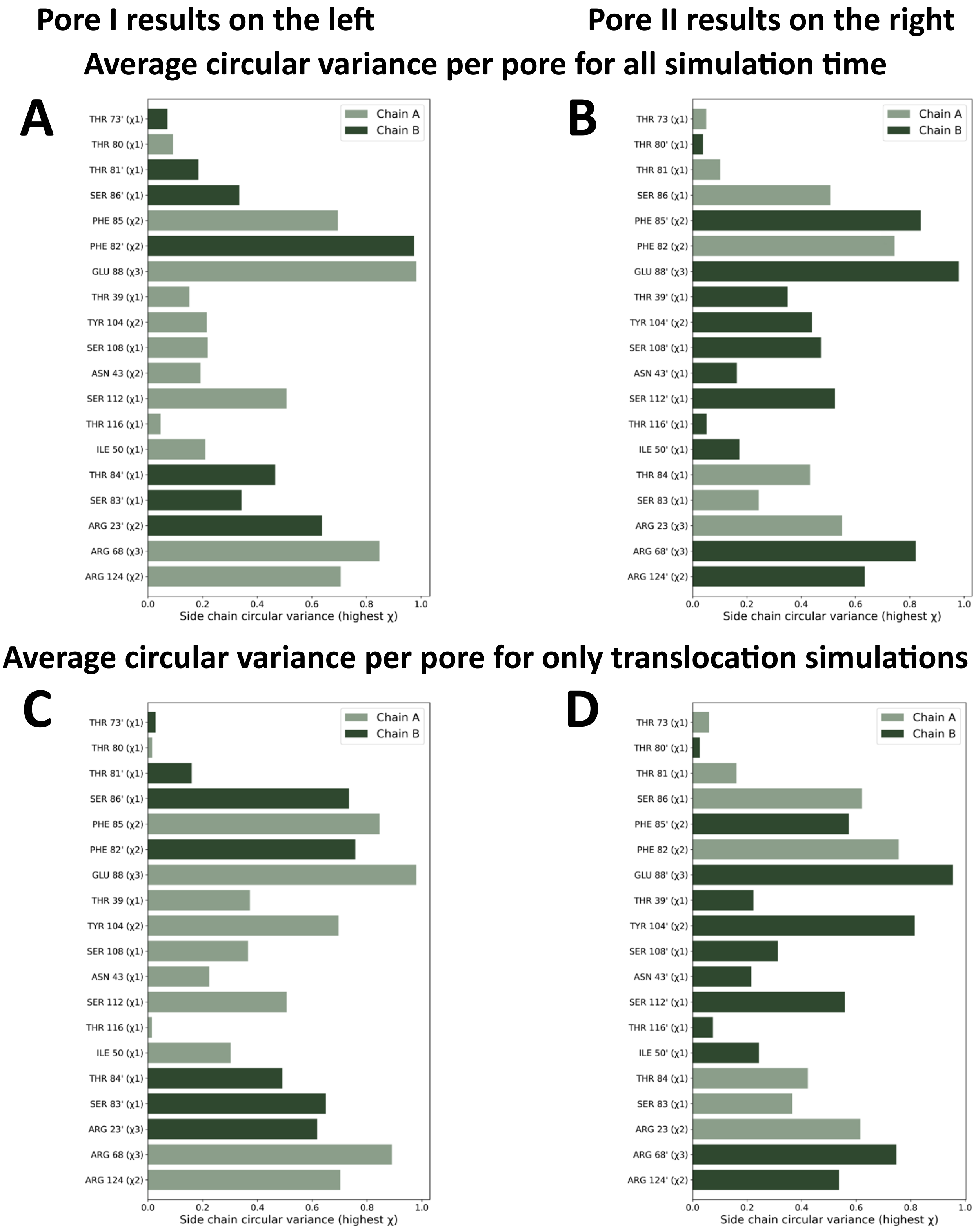


**Fig. S6.** Average dihedral circular variance. (*A*) Average dihedral side chain circular variance for Pore I-lining residues over the entire simulation time. (*B*) Average dihedral side chain circular variance for Pore II-lining residues over the entire simulation time. (*C*) Average dihedral side chain circular variance for Pore I-lining residues calculated only from simulations that exhibited fluoride translocation. (*D*) Average dihedral side chain circular variance for Pore II-lining residues calculated only from simulations that exhibited fluoride translocation.

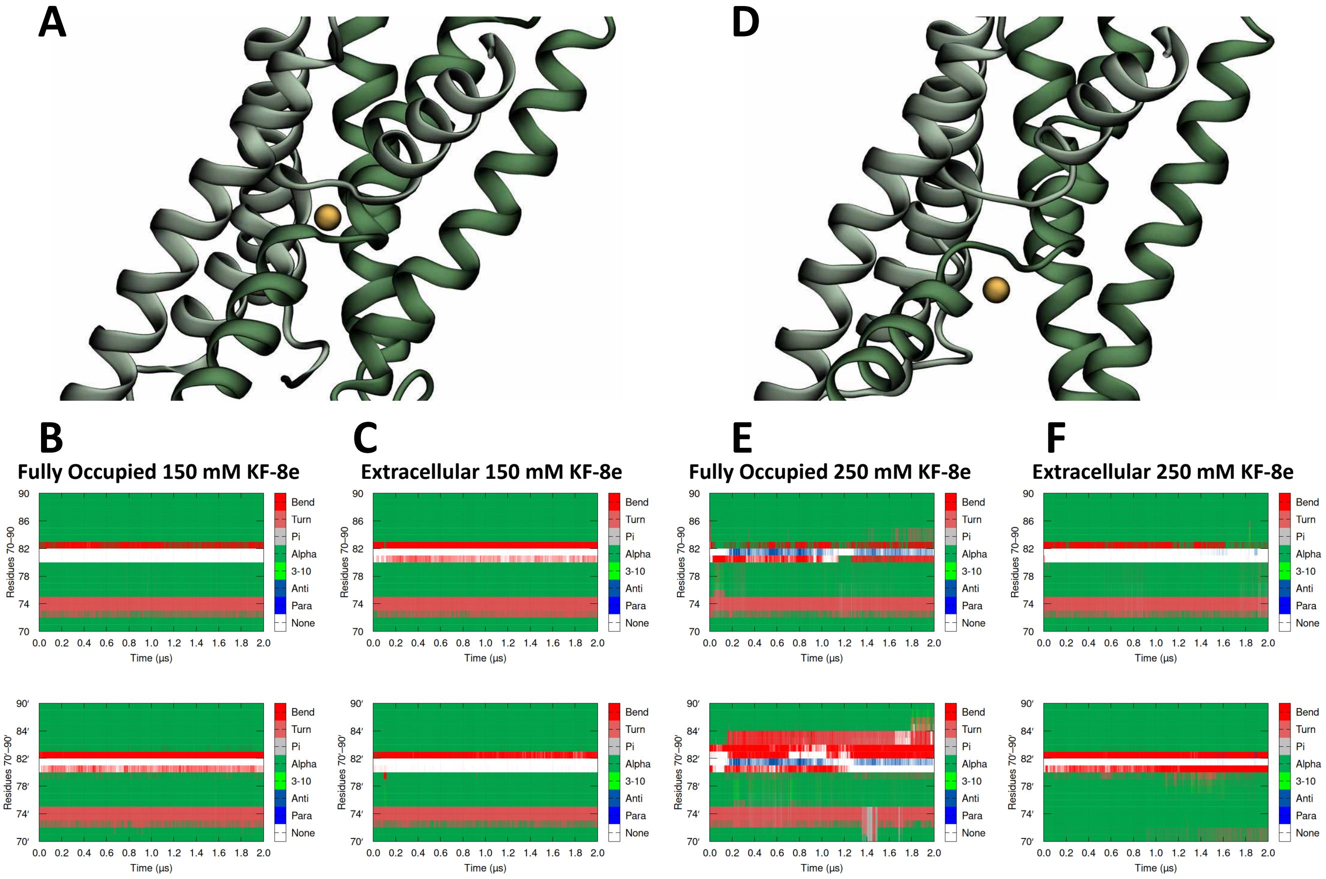


**Fig. S7.** Sodium release from the central cation site in Fluc. (*A*) Representative structural snapshot from a simulation illustrating the typical secondary structure organization around Fluc's central cation site. (*B*) Time-resolved secondary structure analysis for the representative Fully occupied 150 mM KF-8e simulation. (*C*) Time-resolved secondary structure analysis for the representative Extracellular 150 mM KF-8e simulation. (*D*) Representative frame from the Fully occupied 250 mM KF-8e simulation showing atypical secondary structure alterations that may have facilitated sodium release from the central cation site. (*E*) Time-resolved secondary structure analysis for the Fully occupied 250 mM KF-8e simulation, in which sodium dissociation from the central cation site occurred. (*F*) Time-resolved secondary structure analysis for the Extracellular 250 mM KF-8e simulation, which also suffered sodium escape from the central cation site. Transmembrane helix 4′ has been omitted accordingly from the graphical representation for ease of visualization of the ion.

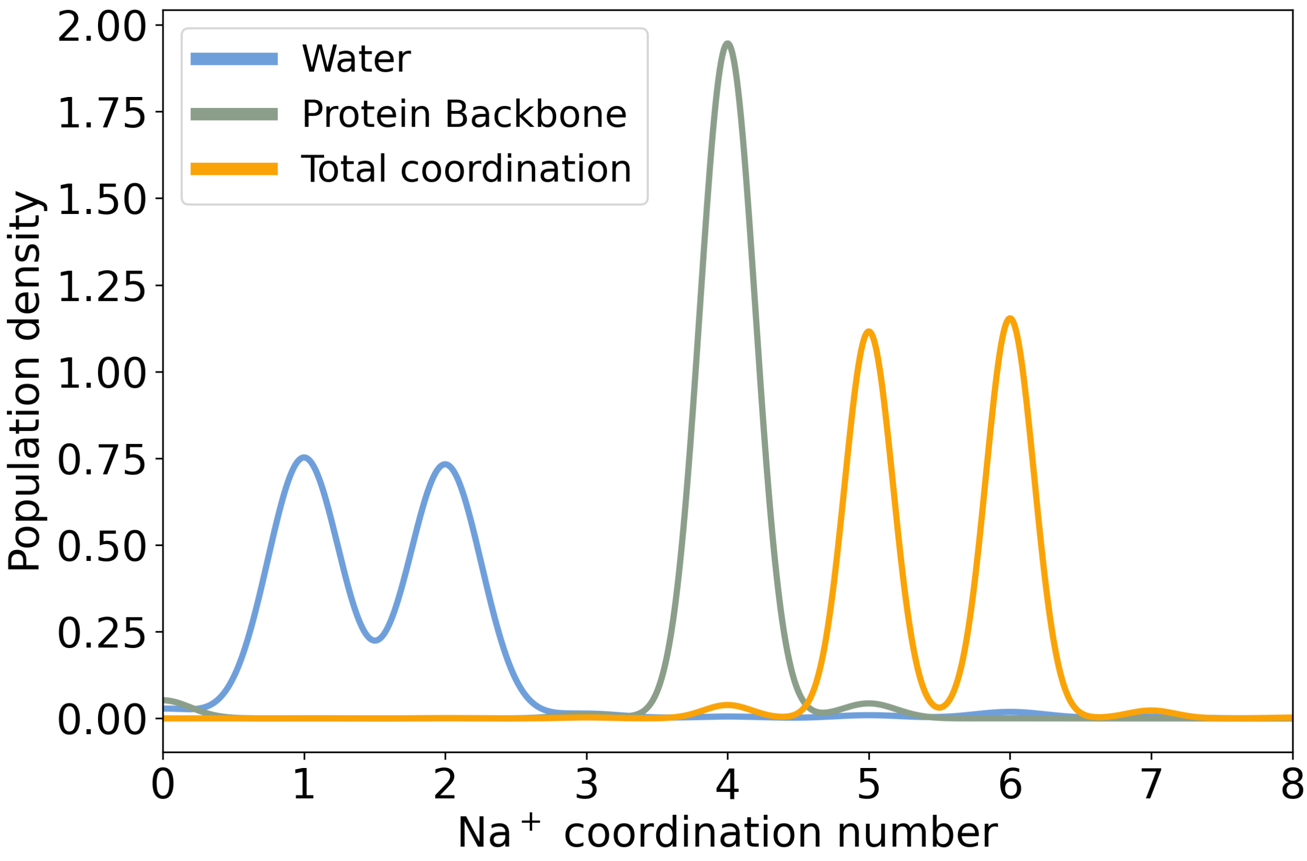


**Fig. S8.** Central sodium coordination across all trials. Total coordination of the central sodium (orange) is shown as the sum of protein carbonyl backbone contacts within 3 $\AA$ (green) and coordinating water molecules (blue), also within 3 $\AA$ of the ion.

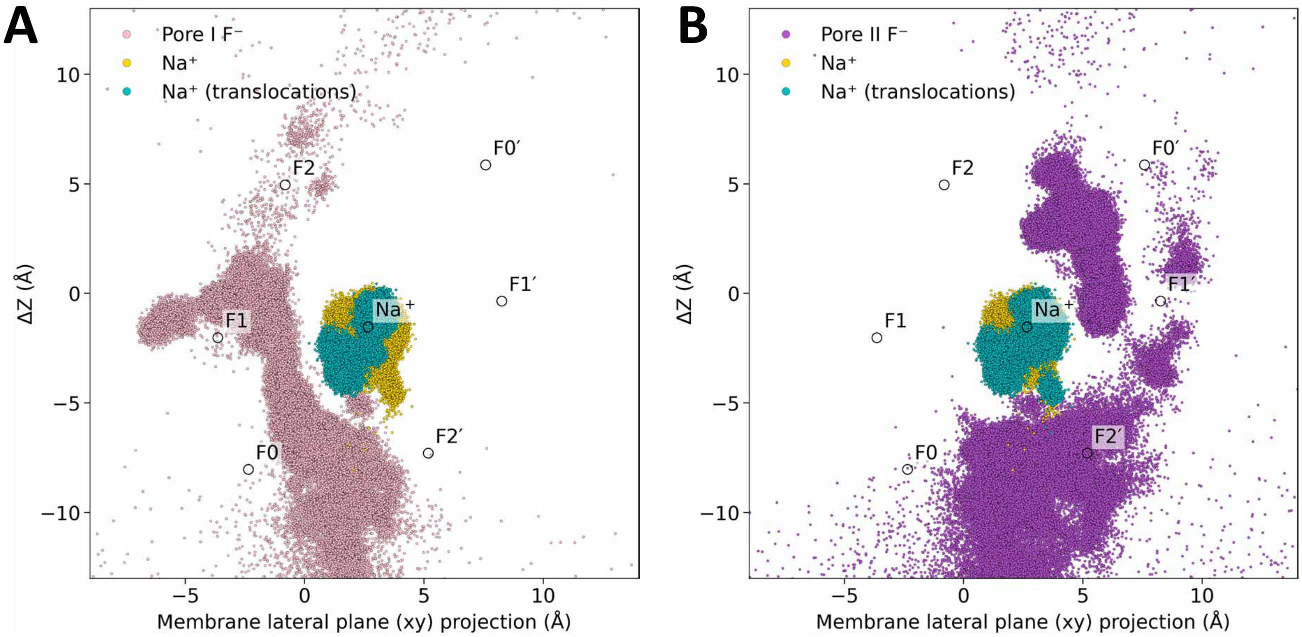


**Fig. S9.** Highlighted central sodium movement during fluoride translocations. (*A*) Scatter plot showing the motion of the central sodium ion (yellow), the sodium positions sampled during trials that exhibited Pore I translocation events (cyan), and Pore I-assigned fluoride ions (pink). (*B*) Scatter plot showing the motion of the central sodium ion (yellow), the sodium positions sampled during trials that exhibited Pore II translocation events (cyan), and Pore II-assigned fluoride ions (pink).

 

**Table S1. Summary of solution charges in Fluc system setup, showing all nine KF concentration and charge difference systems. Each cell shows the total number of cations and anions in both the intracellular (Intra) and extracellular (Extra) solutions, displayed as the number of cations, the number of anions (total charge of the solution), and the total charge of the solution. Additionally, the last row of each concentration reports the membrane potential (△V) achieved for each charge imbalance in V units.**

| Concentration | △q = 0e | △q = 4e | △q = 8e |
|---|---|---|---|
| **150 mM KF** | Intra: 180+, 180– (0)<br>Extra: 83+, 83– (0)<br>△V: 0.158 V | Intra: 178+, 180– (–2)<br>Extra: 85+, 83– (+2)<br>△V: 0.626 V | Intra: 176+, 180– (–4)<br>Extra: 87+, 83– (+4)<br>△V: 1.130 V |
| **250 mM KF** | Intra: 260+, 260– (0)<br>Extra: 83+, 83– (0)<br>△V: 0.339 V | Intra: 258+, 260– (–2)<br>Extra: 85+, 83– (+2)<br>△V: 0.862 V | Intra: 256+, 260– (–4)<br>Extra: 87+, 83– (+4)<br>△V: 1.185V |
| **500 mM KF** | Intra: 390+, 390– (0)<br>Extra: 83+, 83– (0)<br>△V: 0.358 V | Intra: 388+, 390– (–2)<br>Extra: 85+, 83– (+2)<br>△V: 0.872 V | Intra: 386+, 390– (–4)<br>Extra: 87+, 83– (+4)<br>△V: 1.293 V |

**Table S2. Global, pore-specific, and region-partitioned occupancy for anions across:**

| | | All 150 mM KF simulations (18 trials) | | All 150 mM KF & Empty binding site simulations (3 trials) | |
|---|---|---|---|---|---|
| Occupancy type | Region/State | $F^{-}$ (%) | $Cl^{-}$ (%) | $F^{-}$ (%) | $Cl^{-}$ (%) |
| **Global occupancy** | | | | | |
| | None occupied | 12.43 | 63.19 | 29.34 | 49.68 |
| | Pore I only | 50.95 | 21.17 | 67.60 | 34.07 |
| | Pore II only | 31.92 | 13.88 | 1.00 | 13.86 |
| | Both pores | 4.70 | 1.76 | 2.06 | 2.39 |
| **Pore I occupancy** | | | | | |
| | Total | 55.65 | 22.93 | 69.66 | 36.46 |
| | Lower region only (F0–F1) | 49.56 | 18.16 | 69.66 | 35.17 |
| | Upper region only (F2) | 5.24 | 4.56 | 0.00 | 1.02 |
| | Both regions (F0–F1 & F2) | 0.85 | 0.21 | 0.00 | 0.27 |
| **Pore II occupancy** | | | | | |
| | Total | 36.62 | 15.64 | 3.06 | 16.25 |
| | Lower region only (F2′) | 8.81 | 11.58 | 3.06 | 4.51 |
| | Upper region only (F0′-F1′) | 26.94 | 3.87 | 0.00 | 11.29 |
| | Both regions (F2′ & F0′–F1′) | 0.87 | 0.19 | 0.00 | 0.45 |

**Movie S1. Fluoride recruitment and distribution to the intracellular electropositive vestibule.**

**Movie S2. Single fluoride translocation event occurring in Pore I following the proposed Channsporter mechanism.**

**Movie S3. Paired fluoride translocation events occurring in Pore II following the proposed Multi-ion mechanism.**
